# A bioactive phlebovirus-like envelope protein in a hookworm endogenous retrovirus

**DOI:** 10.1101/2021.11.23.469668

**Authors:** Monique Merchant, Carlos P. Mata, Yangci Liu, Haoming Zhai, Anna V. Protasio, Yorgo Modis

## Abstract

Endogenous retroviruses (ERVs), accounting for 14% of our genome, serve as a genetic reservoir from which new genes can emerge. Nematode ERVs are particularly diverse and informative of retrovirus evolution. We identify Atlas virus – a novel, intact ERV in the human hookworm *Ancylostoma ceylanicum* with an envelope protein genetically related to G_N_-G_C_ glycoproteins from phleboviruses. A cryo-EM structure of Atlas G_C_ reveals a class II viral membrane fusion protein fold not previously seen in retroviruses. Atlas G_C_ has the structural hallmarks of an active fusogen. Atlas G_C_ trimers insert into membranes with endosomal lipid compositions and low pH. When expressed on the plasma membrane, Atlas G_C_ has cell-cell fusion activity. Atlas virus is transcriptionally active at specific stages of hookworm development. The preserved activities and expression pattern of Atlas G_C_ suggest it has acquired a cellular function. Our work reveals an unexpected degree of structural and genetic plasticity in retroviruses.

## Introduction

Retroviruses can integrate their genome, reverse-transcribed from RNA into DNA, into the host-cell genome. Viral genomes integrated into germline cells are inherited by future generations as endogenous retroviruses (ERVs). ERVs account for approximately 14% of the human genome, seven times more than protein-coding genes. ERVs and other transposons were initially viewed as parasitic DNA. It is now evident that they serve as a genetic reservoir, from which new genes and regulatory elements can emerge. Sequences of retroviral origin help control gene expression by serving as promoters, enhancers and other regulatory elements (*1, 2*). Genes coopted from ERVs have evolved to fulfill vital cellular functions (*3*). For example, syncytins, which drive cell-cell fusion of trophoblasts during placental development, are encoded by ERV envelope glycoprotein (*env*) genes (*1, 4*). Another recent example is the Gag capsid protein encoded by the *Caenorhabditis elegans* Cer1 retrotransposon, which encapsidates small non-coding RNAs into nucleocapsids that can transfer RNAs conferring learned pathogen avoidance behavior from infected parents to naïve progeny (*5*).

The reduced mutation rate of host versus retrovirus genomes (10^-9^ vs. 10^-3^ mutations per site per year (*6*)) means that ERVs are windows to ancient retroviral sequences – evolutionary fossils preserved from the time of integration. Some ERV genes are expressed in human tissues and retain their biological activities, such as membrane fusion activity in the case of Env proteins (Envs) (*1, 4, 7*). Aberrant expression of Envs is associated with disease (*8, 9*). With the biology of ERVs still largely uncharted, it is likely that many cellular functions of ERVs in health and disease remain undiscovered. Studying ERV genes with novel properties could therefore provide insights on the evolutionary history of retroviruses and identify fundamental principles in host-virus coevolution.

Nematode ERVs are particularly diverse and informative of retrovirus evolution. ERVs from the *Belpaoviridae* (BEL/Pao) family (*10*), widespread across metazoa, have revealing genetic features in nematodes (*11*). The presence in *C. elegans* ERVs of overlapping ORFs, otherwise unique to complex vertebrate retroviruses, suggests retroviruses originated in early metazoa with a common ancestor resembling belpaoviruses (*12*). Furthermore, nematode belpaovirus ERVs encode Envs that are genetically unrelated to all other retrovirus Envs(*13*). Instead of a class I viral membrane fusion protein with a core fold of three bundled α-helices (*14–17*) – a defining feature of modern retroviruses (*18*) – belpaovirus Envs have sequence similarity to G_C_ (G2) envelope glycoproteins from phleboviruses (family *Phenuiviridae*) (*13*). G_C_ proteins are class II membrane fusion proteins, with a three-domain β-strand architecture (*19, 20*) also found in alphaviruses (*21*), flaviviruses (*22, 23*) and Rubella virus (*24*) but structurally unrelated to class I fusion proteins. A series of conformational changes in class II fusion proteins, triggered by endosomal acidification, catalyses fusion of the viral and endosomal membranes to deliver the viral genome into the cytosol (*25–27*). A hydrophobic fusion loop first inserts into the endosomal membrane. The proteins then form trimers and fold back on themselves, pulling the cell membrane (held by the fusion loop) and the viral membrane (held by a transmembrane anchor) together so they fuse (*18, 20*).

Class II fusion proteins are not limited to viruses: they also drive cell-cell fusion events of fundamental importance, including syncytial epithelia formation in *C. elegans* (*28*), and gamete fusion in protozoa, plants, algae and invertebrates (*29–31*). The identical topology and overall arrangement of the three domains of viral and eukaryotic class II fusion proteins, along with similarities in their membrane fusion mechanisms, makes it all but certain they evolved from a common ancestor (*29*). Although the evolutionary origin of the ancestral class II fusion protein remains unknown, the presence of class II fusion proteins in ERVs raises the provocative prospect that a gene transfer from a virus to a cell led to the advent of sexual reproduction (*29, 32*).

Here, we identify a novel, intact belpaovirus ERV in the human hookworm *Ancylostoma ceylanicum* (a parasitic nematode) with an Env more similar than any other eukaryotic sequence to phlebovirus G_C_ protein sequences. We expressed and purified the G_C_-homologous fragment from this ERV, henceforth Atlas virus. A cryo-electron microscopy (cryo-EM) structure of Atlas G_C_ reveals a class II viral fusion protein fold remarkably similar to phlebovirus G_C_ proteins and not previously seen in retroviruses. We show that Atlas G_C_ has all the hallmarks of an active class II membrane fusion protein. It undergoes a monomer-to-trimer transition and inserts into lipid membranes with a specific lipid composition in response to a low pH trigger. Our work provides biochemical validation for the hypothesis that acquisition of a fusion protein from an infectious virus, as exemplified by Atlas virus, represents a general paradigm for how retrotransposons can become retroviruses (*13*), and indeed how ancestral reverse-transcribing viruses may have originated (*10*). The preserved biological activities of Atlas G_C_, including membrane fusion activity, raise the question of whether these activities, and those of ERV gene products more broadly, have cellular functions or cause disease.

## Results

### An intact ERV with a phlebovirus-like Env in the hookworm *A. ceylanicum*

The bioinformatic discovery of nematode ERVs with phlebovirus G_C_-like Env sequences not previously seen in retroviruses (*11–13*), requires biochemical validation. To identify phlebovirus-like ERV Envs suitable for biochemical analysis, we performed PSI-BLAST searches (*33*) for protein sequences similar to biochemically characterized phlebovirus G_C_ proteins. A search with Rift Valley Fever virus (RVFV) G_C_ as the query identified the gene *Acey_s0020.g108* (UniProt A0A016UZK2) in the human hookworm *A. ceylanicum* as containing the most similar sequence outside infectious virus taxa (Expected value 10^-20^). The homologous sequence lies within a 9,204-nucleotide element possessing all the features of an intact ERV, including 100% identical 271-nt LTRs and a coding sequence encoding a single 2,828-residue Gag-Pol-Env polyprotein without any stop codons or introns (**Fig. 1a; Supplementary Fig. 1a**). We refer to this element as *A. ceylanicum* Atlas virus. It is one of nine *A. ceylanicum* ERVs that encode complete Gag-Pol-Env polyproteins (**Fig. 1b,c**). These ERVs have the distinguishing genomic features of belpaoviruses from other nematode species, including an atypical Env and an aspartate to asparagine substitution (Y[X]D<u>D</u> → YVD<u>N</u>) in the most conserved reverse transcriptase motif, the polymerase site (*12, 34, 35*) (**Supplementary Fig. 1b**). With a Gag sequence less than 20% identical to its closest homolog (*Acey_s0020.g1106*), Atlas virus is a candidate for classification as a member of the *Belpaoviridae* family (which contains a single genus, *Semotivirus*) (**Supplementary Fig. 1c**).

**Figure 1.**
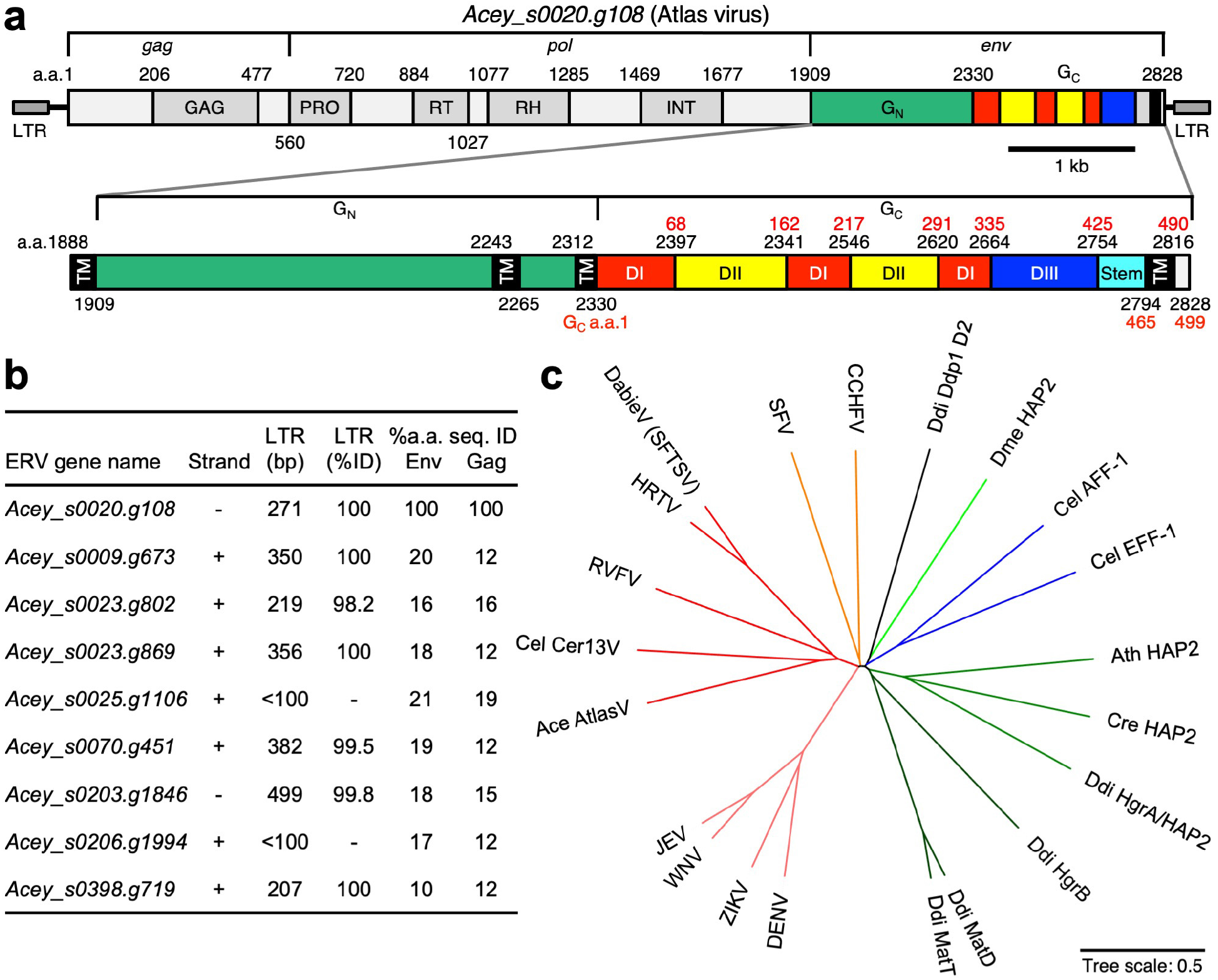
Phylogenetic analysis of class II-like Env genes and their closest relatives from modern infectious viruses. **(a)** Gene architecture of *Acey_s0020.g108*, an endogenous retrovirus with a phlebovirus-like Env from the *Ancylostoma ceylanicum* human hookworm parasite. We refer to this element here as *A. ceylanicum* Atlas virus. Inset: annotated closeup of the Atlas Env region encoding phlebovirus G_N_- and G_C_-like glycoproteins. Red residue number refer to the G_C_ sequence alone. **(b)** List of *A. ceylanicum* ERVs encoding complete Gag-Pol-Env polyproteins with phlebovirus- like Envs. The length and sequence identity of LTRs is listed. LTRs sequence identities for two of the genes could not be calculated: *Acey_s0206.g1994* had degenerate LTRs, and one of the LTRs of *Acey_s0025.g1106* was truncated in the genome assembly. Amino acid sequence identities of Env and Gag sequences are listed. **(c)** Phylogenetic tree of G_C_ proteins from *Phenuiviridae* and G_C_-like (G2-like) sequences from retrotransposons. Ace, *A.* ceylanicum; Cel, *C. elegans*; Ath, *Arabidopsis thaliana*; Cre, *Chlamydomonas reinhardtii*; Ddi, *Dictyostelium discoideum*; CCHFV, Crimean Congo hemorrhagic fever virus; SFV, Semliki Forest virus; DabieV, Dabie bandavirus; HRTV, Heartland virus; JEV, Japanese encephalitis virus; WNV, West Nile virus; ZIKV, Zika virus; DENV, dengue virus. Drawn with Interactive Tree Of Life (iTOL) (*36*).

The phlebovirus G_C_-like sequence spans the last 498 residues of the Atlas virus polyprotein (residues 2330-2828). It contains a single predicted C-terminal transmembrane helix, like phlebovirus G_C_ proteins. Phleboviruses and other *Phenuiviridae* family members express a glycoprotein precursor that is cleaved by cellular proteases into two envelope glycoproteins, G_N_ and G_C_ (or G1 and G2). G_N_ is the receptor-binding protein required for cellular attachment and G_C_ is the membrane fusion protein required for cell entry. G_N_ is highly antigenic and more variable in sequence than G_C_. Our analysis of the Atlas virus Env sequence detected a slight but statistically significant similarity in the 421 residues preceding the G_C_-like sequence (residues 1909-2329) to G_N_ glycoproteins from *Phenuiviridae* family viruses (*E*-value *≥* 10^-7^). Moreover, the distribution of predicted transmembrane helices and proteolytic cleavage sites in and adjacent to the Atlas G_N_- and G_C_-like sequences is the same as in phlebovirus glycoproteins (**Fig. 1a**). Together, these sequence features suggest the Atlas virus Env contains tandem phlebovirus-like G_N_ and G_C_ glycoproteins instead of a retrovirus-like glycoprotein. With all the features of a recently active ERV and an apparently intact set of phlebovirus-like glycoproteins, Atlas virus is an excellent candidate for biochemical analysis.

### Atlas G_C_ has a class II membrane fusion protein fold not previously seen in retroviruses

As the molecular structure of phlebovirus G_C_ proteins, and how they drive fusion of the viral and cellular membranes, are well established from previous studies (*19, 27*), we focused our biochemical analyses on the G_C_-like sequence from Atlas virus. A recombinant ectodomain fragment of Atlas G_C_ (polyprotein residues 2330-2772) was expressed in *Drosophila melanogaster* D.mel-2 cells as a secreted protein. The purified protein was a soluble, folded homotrimer (**Supplementary Fig. 2**). The structure of the G_C_ trimer was determined by single-particle cryo-EM image reconstruction at an overall resolution of 3.76 Å (**Fig. 2a,b; Supplementary Table 1; Supplementary Fig. 3**). The map was sufficiently detailed for an atomic model to be built and refined for Atlas G_C_ residues 2330-2769 using the crystal structure of RVFV G_C_ (*27*) as a starting model (see Methods; **Supplementary Fig. 4**). The structure reveals a three-domain class II membrane fusion protein fold (**Fig. 2c**). All previously described retroviral Env structures have a helical coiled-coil class I fusion protein fold (*14–17*), exemplified by the hemagglutinin of influenza virus (*18*). The structure of the Atlas G_C_ ectodomain fragment is remarkably similar to phlebovirus G_C_ structures, specifically the trimeric postfusion G_C_ structures from RVFV (*27*), Dabie bandavirus (DABV, formerly SFTS phlebovirus) (*37*) and Heartland virus (HRTV) (*38*) (**Fig. 2d-e**). Domain I, a 10-stranded *β*-barrel augmented by a 3-stranded sheet, organizes the structure. Two insertions in domain I form the elongated, mostly *β*-stranded domain II. Domain III has the 7-stranded *β*-sandwich topology of fibronectin type III (FN3) domains also found in macroglobulin domains (*31, 39*), but the hydrophobic core and disulfide bonding pattern of domain III differ from these and other annotated domains from non-viral species. A 16-amino acid portion of the stem region, which links domain III to the C-terminal transmembrane anchor in class II fusion proteins, could be modeled, spanning 5 nm from the end of domain III to within approximately 1 nm of the tip of domain II. The stem forms trimer contacts, adding a *β*-strand to domain II of a different subunit, as seen in RVFV G_C(27)_. The overall configuration bears strong similarity to other viral and cellular class II fusion proteins including, in order of decreasing similarity: alphavirus E1 proteins (*21*); EFF-1/AFF-1 cell-cell fusion proteins from *C. elegans* (*28*); HAP2 gamete fusion proteins from protozoa (*29–31, 40*) and plants (*30*); and flavivirus E proteins (*41*) (**Fig. 2e**). Despite these structural similarities, the only proteins or domains of known structure with detectable amino acid sequence similarity to Atlas G_C_ (*E*-value <1 in PSI-BLAST (*33*)) are the phlebovirus G_C_ proteins (22-24% sequence identity; **Supplementary Fig. 5**).

**Figure 2.**
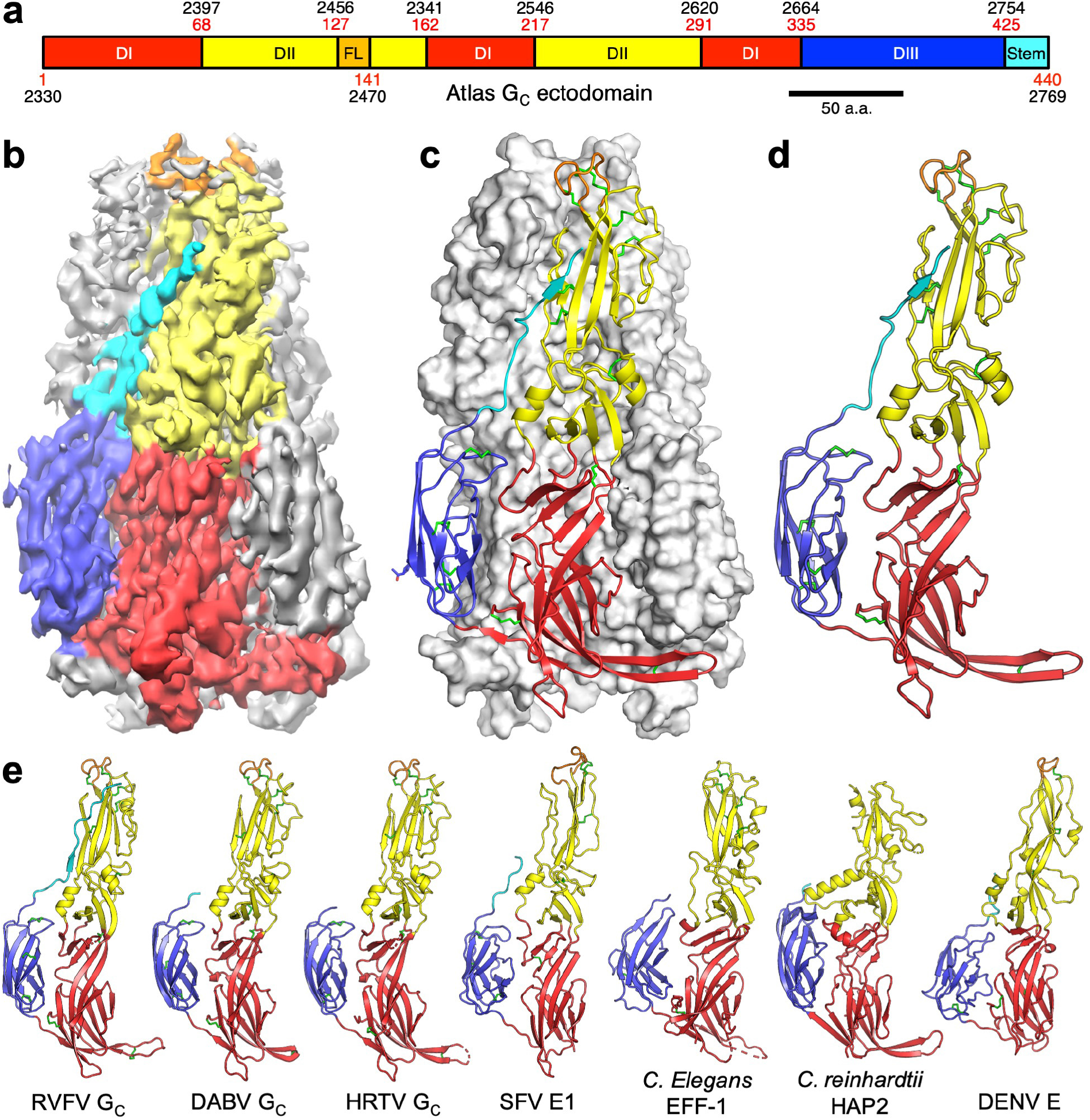
Cryo-EM structure of a phlebovirus G_C_-like Env fragment from *A. ceylanicum* Atlas virus. **(a)** Domain organization of Atlas virus G_C_. Domains are colored as in Fig. 1a. **(b)** 3D density map of the cryo-EM image reconstruction of a soluble ectodomain fragment of Atlas virus G_C_ at 3.76 Å overall resolution. The purified protein was a homotrimer (**Supplementary Fig. 2**) and threefold (C3) symmetry was imposed. The map is colored by domain as in **(a).** A representative cryo-EM micrograph is shown in **Supplementary Fig. 4a**. **(c)** Overview of the refined atomic model of the Atlas virus G_C_ trimer. Atlas G_C_ has the fold of a class II fusion protein in its postfusion conformation. The protomer in the foreground is shown as a cartoon and the two other protomers in surface representation. Disulfide bonds (green) and an N-linked glycan (blue) are shown as sticks. **(d)** Protein fold of single Atlas G_C_ protomer. **(e)** The structures most similar to Atlas G_C_ are G_C_ glycoproteins from Rift Valley Fever virus (RVFV, Rmsd 2.6 Å, Z-score (*42*) 29, PDB:6EGU (*27*)), Dabie bandavirus (DABV, formerly SFTS phlebovirus; Rmsd 2.6 Å, Z-score 28, PDB:5G47 (*37*)) and Heartland virus (HRTV, Rmsd 2.6 Å, Z-score 28, PDB:5YOW (*38*)). Other representative class II fusion proteins are shown for comparison: Semliki Forest virus (SFV) E1 (Z-score 19, PDB:1RER (*26*)), *C. elegans* EFF-1 (Z-score 19, PDB:4OJD (*28*)), *Chlamydomonas reinhardtii* HAP2 (Z-score 16; PDB:5MF1 (*29*)) and dengue virus (DENV) E (Z-score 15, PDB:4GSX (*43*)).

Atlas G_C_ has the same structural features that distinguish phlebovirus glycoproteins from other class II fusion proteins: a larger number of disulfide bonds, ten of which are conserved in phlebovirus G_C_ sequences but not in other class II proteins; N-linked glycosylation in domain III; and a more extensive and rigid interface between domains I and II (**Fig. 2d-e; Supplementary Fig. 5**). The most notable differences between Atlas virus and phlebovirus G_C_ structures are differences in the disulfide bonding pattern and in the composition of side chains lining the glycerophospholipid (GPL) headgroup binding pocket conserved in arboviral class II fusion proteins (*27*). We discuss these differences and their potential functional implications below. Atlas G_C_ also has a different glycosylation pattern, with a single predicted N-linked glycosylation site at Asn414 in domain III with a weak corresponding feature in the density map (**Supplementary Fig. 6**). In contrast, phlebovirus G_C_ proteins contain two N-linked glycans in domain III, at two different sites. One of these, Asn1035 in RVFV, covers the fusion loop in the prefusion conformation of RVFV G_C_ and stabilizes the prefusion dimer by forming contacts across the dimer interface (*19*), as also seen in flavivirus E proteins (*22, 23*). This glycosylation site is conserved in DABV and HRTV G_C_ but absent in Atlas G_C_. Despite these minor differences, the striking overall structural similarity of Atlas G_C_ to phlebovirus G_C_ proteins in the postfusion conformation reveals a surprisingly strong evolutionary link between nematode ERVs from the *Belpaoviridae* family and the fusion proteins of phleboviruses.

### Structure of the putative lipid membrane anchor of Atlas virus G_C_

Viral fusion proteins insert a membrane anchor – the fusion loop, in class II proteins – into the host cell membrane to initiate virus-cell membrane fusion. The putative fusion loop of Atlas G_C_ can be clearly identified by analogy to phlebovirus G_C_ proteins as spanning residues 127-140. The local resolution of the cryo-EM density for this region is lower than for the rest of the map but the large number of structural constraints imposed by the positions of disulfide-bonded cysteines and other residues conserved in phleboviruses G_C_ proteins allowed an atomic model to be built unambiguously (**Fig. 3a; Supplementary Fig. 6b,c**). Specifically, the structure of the fusion loop is constrained by the positions of four disulfide bonds conserved in phleboviruses, a fifth disulfide specific to Atlas G_C_ (Cys129-Cys138), a phenylalanine (Phe136) required in phleboviruses at the apex of the fusion loop for membrane binding and fusion (*27, 37, 44*), and two conserved glycines (Gly128 and Gly134) that provide the torsional flexibility necessary for the fusion loop’s tightly folded conformation (**Fig. 3a-c**).

**Figure 3.**
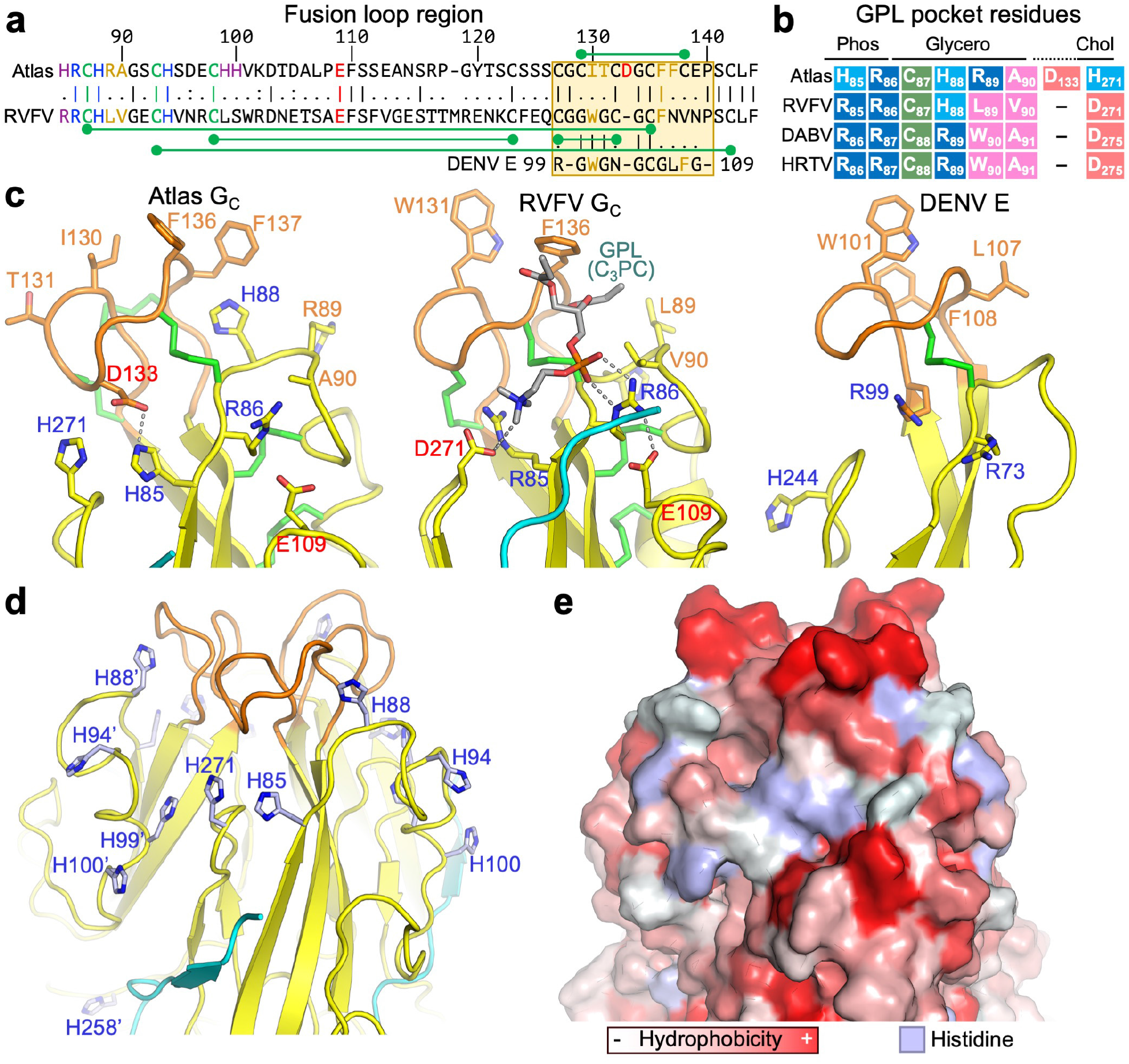
Structure of the putative fusion loop and lipid binding pocket of Atlas G_C_. **(a)** Local alignment of the sequences surrounding the fusion loops of Atlas G_C_, RVFV G_C_, and DENV E. Fusion loop sequences are boxed in orange. Disulfides are marked with green dumbbells. Disulfides shown below the RVFV sequence are conserved in Atlas and RVFV. **(b)** Residues contributing to binding of the phosphate (Phos), glycerol and choline (Chol) moieties of glycerophospholipid (GPL) molecules by RVFV G_C_ (*27*). The aligned sequences of G_C_ from Atlas, Dabie bandavirus (DABV, formerly SFTSV) and heartland virus (HRTV) are shown. Colors indicate side chain properties: blue, positive charge; pink, negative charge; magenta, hydrophobic; green, sulfhydryl; light blue, positive charge at endosomal pH. **(c)** Closeup of the Atlas G_C_ fusion loop and GPL binding pocket structure. The corresponding structures of RVFV G_C_ and dengue E are shown for comparison. A single protomer from the postfusion trimer is shown, with key residues shown in stick representation. Label colors indicate side chain properties: blue positive charge; red, negative charge; orange, hydrophobic. **(d)** Closeup of the fusion loop and GPL binding pocket region of the Atlas G_C_ trimer showing histidine residues in or near the pocket. Prime symbols following residue numbers denote the protomer to which the residue belongs. **(e)** Surface representation of the same view as in (**d**), colored by side chain hydrophobicity, except for histidine residues shown in light blue.

The structure and chemical properties of the Atlas G_C_ fusion loop are similar to phlebovirus fusion loops and consistent with a membrane anchoring function. The side chains of Ile130, Phe136, Phe137 and the Atlas-specific disulfide (Cys129-Cys138) form a hydrophobic surface at the narrow end of the trimer similar to phlebovirus and flavivirus fusion proteins (*25, 27*) (**Fig. 3c**). The area of this surface, extended by the side chain of Arg89 on an adjacent loop, analogous to Leu779 in RVFV G_C_, is greater than in most other viral class II fusion proteins. By analogy with other viral class II fusion proteins, the location and extent of the hydrophobic surface formed by the Atlas G_C_ fusion loop suggest it could function as a membrane anchor.

In addition to inserting nonpolar side chains into the hydrophobic region of the membrane, viral class II fusion proteins form polar contacts with lipid headgroups via the fusion loop and an adjacent glycerophospholipid (GPL) binding pocket (*27*). By selecting for headgroups with complementary electrostatic potential, polar contacts confer a degree of specificity to lipid binding. In phleboviruses, a set of conserved polar residues in the GPL binding pocket bind selectively to zwitterionic GPLs (*27*). The Atlas G_C_ structure reveals a putative GPL binding pocket with both conserved and novel features (**Fig. 3**). The arginine that forms bidentate hydrogen bonds with the GPL phosphate moiety in phleboviruses (*27*) is conserved in Atlas virus (Arg86). The disulfide bond and short-chain hydrophobic residue that bind the GPL glycerol moiety are also conserved (Cys87-Cys135, Ala90). However, an aspartate-arginine pair that binds choline and ethanolamine GPL moieties in phleboviruses is not conserved in Atlas G_C_, which instead has two histidines at the corresponding positions (His271 and His85). Moreover, Atlas virus has an extra residue in the fusion loop, Asp133, compared to phlebovirus G_C_ proteins. The side chain of Asp133 points into the GPL binding pocket and is located near the position of the choline GPL moiety in the superimposed structure of RVFV G_C_ bound to a phosphatidylcholine ligand (*27*), suggesting that Asp133 could compensate for the lack of a conserved aspartate at position 271 (**Fig. 3a-c**). Hence the GPL binding pocket of Atlas G_C_ appears to have the necessary physicochemical attributes to support GPL binding, with Arg86 binding the phosphate moiety, Cys87/Cys135/Ala90 binding the glycerol moiety, and His85/Asp133/His271 coordinating the end of the headgroup. We note the presence in the cryo-EM reconstruction of a bulge in the density around the GPL binding pocket that is unaccounted for by the atomic model (**Supplementary Fig. 6c**). Additionally, the absorbance at 260 nm of purified Atlas G_C_ was higher than expected despite treatment with nucleases during purification (**Supplementary Fig. 2a**). These two observations would be consistent with lipid molecules with unsaturated acyl chains copurifying with Atlas G_C_, but the local resolution of the map is insufficient to ascertain whether the GPL binding pocket contains a ligand.

### Atlas G_C_ binds membranes with endosome-like lipid composition at low pH

A key step in viral membrane fusion is insertion of the fusion protein into the host cell membrane. To determine whether Atlas G_C_ has this activity, we assessed binding to liposomes in density gradient centrifugation experiments, supported by dynamic light scattering (DLS) measurements. Viruses containing class II fusion proteins, like many retroviruses, undergo membrane insertion and fusion in endosomal compartments where the pH is acidic. We therefore assayed liposome binding at a range of pH values. In contrast to RVFV G_C_, Atlas G_C_ did not bind liposomes containing phosphatidylcholine (PC), phosphatidylethanolamine (PE), cholesterol and sphingomyelin (SM) at neutral or acidic pH (pH 4-8; **Fig. 4a,b**). At neutral pH (pH 7.8), Atlas G_C_ also failed to bind liposomes containing anionic lipids enriched in early or late endosomes: phosphatidylcholine (PS) or bis(monoacylglycerol)phosphate (BMP, also known as lysobisphosphatidic acid or LBPA), respectively. At pH 4, however, Atlas G_C_ bound tightly to liposomes containing PS or BMP, with weaker binding observed at pH 4.6 (**Fig. 4a,b; Supplementary Fig. 7**). Remarkably, Atlas G_C_ bound only weakly to liposomes containing phosphatidylglycerol (PG) instead of BMP even though PG and BMP are regioisomers with identical chemical composition and electrostatic charge (of -1). PG and BMP differ only in the position of the second acylglycerol linkage, resulting in a linear configuration for BMP instead of the usual branched configuration for PG. Our liposome binding data show that Atlas G_C_ binds to membranes containing specific GPLs that are enriched in the endosomal pathway in a pH-dependent manner. No other class II fusion protein has been reported to require low pH, PS or BMP for membrane insertion. However, phleboviruses, which require only PE or PC and cholesterol for membrane insertion (*27*), subsequently require BMP and low pH for fusion (*45*). Similarly, flaviviruses require BMP, PS or other anionic lipids and low pH for efficient fusion (*46–48*).

**Figure 4.**
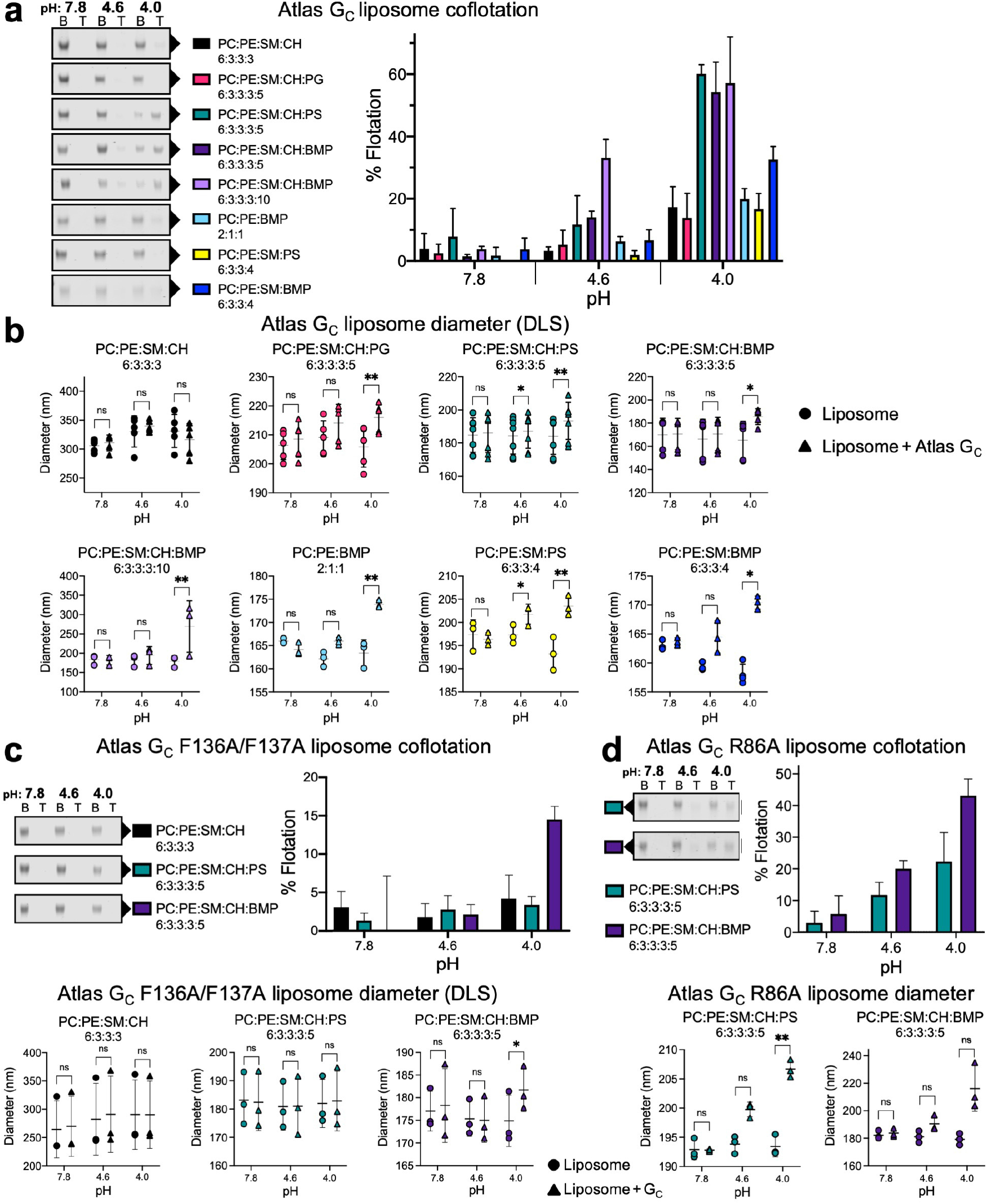
Atlas G_C_ binds liposomes in a lipid- and pH-dependent manner. **(a)** Liposome coflotation lipid binding assay. A liposome-G_C_ mixture in 40% OptiPrep density gradient medium was overlaid with a 30% OptiPrep cushion and centrifuged at 100,000 g. Flotation was defined as the amount of Atlas G_C_ co-floating with liposomes in the top-half fraction (T) divided by the total amount of Atlas G_C_ in the top- and bottom-half (B) fractions. Atlas G_C_ was quantified by Coomassie-stained SDS-PAGE. **(b)** Binding of Atlas G_C_ to liposomes measured by dynamic light scattering (DLS). Liposome diameter was measured by DLS in the presence and absence of Atlas G_C_. **(c)** Binding of Atlas G_C_ F136A/F137A to liposomes measured by flotation and DLS as above. **(d)** Binding of Atlas G_C_ R86A to liposomes measured by flotation and DLS as above. See **Supplementary Dataset 1** for source data and **Supplementary Fig. 8** for uncropped gels for all replicates. Error bars for flotation data show the standard deviation with three replicates except PC:PE:SM:CH:PS + WT (two replicates) and PC:PE:SM:BMP + WT (four replicates); error bars for DLS data show the standard deviation of three to seven replicates. Significance was determined by 2-way ANOVA analysis of the mean change in liposome diameter, using Sidak’s multiple comparisons test with a 95% confidence interval in GraphPad Prism 8 (see **Supplementary Fig. 7**). **, 0.001 < *p* < 0.01; *, *p* < 0.05; ns, not significant.

The optimal pH for membrane insertion of Atlas G_C_ (pH 4-4.5) is similar to the optimal pH of hemifusion of Uukuniemi virus (*45*), a model phlebovirus, and would be consistent with membrane insertion in late endosomes or endolysosomes, as is the case for phleboviruses (*49*). Since the imidazole side chain of histidine has a p*K*_a_ value in the range of 6-6.4, the side chains of His85 and His271, in the GPL binding pocket of Atlas G_C_, would be fully protonated at pH 4-4.5. The resulting net positive charge of the His85/Asp133/His271 triad (+1/-1/+1), analogous to the Arg/Asp pair that coordinates the end of GPL headgroups in phleboviruses, would precisely mirror the charge of the phosphoserine headgroup of PS (−1/+1/-1), and to a lesser extent the glycerol-phospho-glycerol headgroup of BMP (−1). Moreover, Atlas G_C_ contains four additional histidines (residues 88, 94, 99 and 100) exposed to the solvent in the immediate vicinity of the GPL binding pocket and fusion loop (**Fig. 3d,e**), none of which are conserved in phleboviruses. The protonation of these histidines at low pH may promote further interactions with anionic moieties of lipid headgroups, complementing the function of His85/His271. The presence of six histidines in and around the GPL binding pocket provides a possible explanation for the observed pH-dependent insertion of Atlas G_C_ into membranes containing PS and BMP. Consistent with a conserved role for the GPL binding pocket in determining lipid specificity of class II fusion proteins, mutations in alphaviruses at a position equivalent to His271 in Atlas G_C_, in the *ij* loop, determine the extent to which alphaviruses depend on cholesterol for membrane binding (*50*).

### Atlas G_C_ membrane insertion does not strictly require cholesterol and occurs via the fusion loop

In addition to GPLs, phleboviruses and alphaviruses (but not flaviviruses) require cholesterol for efficient membrane binding and subsequent fusion (*27, 45, 50*). Alphaviruses additionally require sphingolipids (such as SM) for efficient fusion (*51, 52*). We found that neither cholesterol nor SM were required for Atlas G_C_ to bind liposomes (**Fig. 4a,b**). Removal of cholesterol reduced the fraction of G_C_ bound by 50% in the liposome flotation assay (**Fig. 4a**) though binding was still detected in the DLS assay (**Fig. 4b**). Hence, although cholesterol and SM are not strictly required for binding, they enhance binding, possibly by increasing membrane fluidity. Notably, the concentration of cholesterol in nematode cell membranes is approximately 20 times lower than in vertebrates (*53, 54*). This is insufficient for cholesterol to regulate the structure or fluidity of nematode membranes, in which cholesterol is thought to be instead a precursor for low-abundance metabolites (*53–55*). Likewise, *Drosophila* can grow indefinitely with only trace amounts of exogenous sterols (*53*), suggesting that arthropods, which are obligate vectors of the vast majority of viruses containing class II fusion proteins, rely on lipids other than cholesterol to regulate membrane fluidity.

In the structure of Atlas G_C_, the hydrophobic residues Ile130 and Phe137 in the fusion loop occupy the positions of more hydrophilic side chains (Trp821 and Asn827 in RVFV G_C_), which are conserved across phleboviruses and flaviviruses and have been implicated in cholesterol binding. Indeed, molecular dynamics simulations suggested that the side chains of Trp821 and Asn827 in the RVFV fusion loop form hydrogen bonds with cholesterol upon membrane insertion (*27*). Moreover, the N827A RVFV G_C_ mutant lacks fusion activity (*44*), and the N827A and W821H mutants both fail to bind membranes (*27*). We speculate that the observed ability of Atlas G_C_ to bind GPL membranes lacking cholesterol – a functionality consistent with the low cholesterol abundance in nematodes – may be due to its more hydrophobic fusion loop surface, with Ile130, Phe137 and the Cys129-Cys138 disulfide complementing the conserved Phe136.

To determine whether Atlas G_C_ binds membranes in a manner analogous to other viral class II fusion proteins – via the fusion loop and GPL binding pocket – we generated Atlas G_C_ variants with mutations in the two phenylalanine residues in the fusion loop (F136A/F137A), or in the arginine predicted to bind the GPL phosphate moiety (R86A). The F136A/F137A mutant failed to bind liposomes containing PS and coflotation with liposomes containing BMP at pH 4 and pH 4.6 was reduced to approximately one third of wild-type (**Fig. 4c**). The R86A mutation reduced binding to liposomes containing PS or BMP at pH 4, to one third and 75% of wild-type, respectively (**Fig. 4d**). For both variants, preparations contained trimers as the major species but a small monomeric fraction was also present (**Supplementary Fig. 2**), suggesting that the mutated lipid binding residues are required for efficient trimer assembly. We conclude that Atlas G_C_ binds to lipid membranes through insertion of hydrophobic fusion-loop residues and coordination of lipid headgroups in the GPL binding pocket, as in other viral class II fusion proteins.

### Evidence for monomeric and trimeric states of Atlas G_C_

Class II fusion protein ectodomains can be monomeric, dimeric or form icosahedral shells in their prefusion conformation, but the fusogenic conformational change is always accompanied by reorganization into trimers (*25, 45, 56, 57*). Fusion proteins from classes I and III, including retrovirus fusogens, remain trimeric throughout the fusion reaction, but no class II fusion proteins are known to be trimeric in their prefusion conformation. Having established that Atlas G_C_ can insert into membranes as a trimer with a postfusion-like conformation, we set out to determine whether it could undergo a conformational change as seen in the fusion reaction of class II proteins from infectious viruses. The Atlas G_C_ construct described above was expressed as a trimer with no trace of monomers or dimers (**Fig. 5a, Supplementary Fig. 2a**). However, we found that a construct with the stem region truncated, G_C_(DI-III), spanning residues 2330-2751, was expressed as a mixture of monomers, trimers and higher order oligomers (**Fig. 5b, Supplementary Fig. 2b**). In contrast to G_C_ trimers, which were stable at different protein concentrations and pH values, G_C_(DI-III) monomers were unstable over time. As noted above, monomeric fractions were also present in preparations of the fusion loop mutant (F136A/F137A) and GPL binding pocket mutant (R86A) (**Supplementary Fig. 2c,d**). Whether these monomeric species are in a prefusion conformation remains to be determined, but the presence of metastable monomers and stable trimers recapitulates a key property of class II fusogens from infectious viruses.

**Figure 5.**
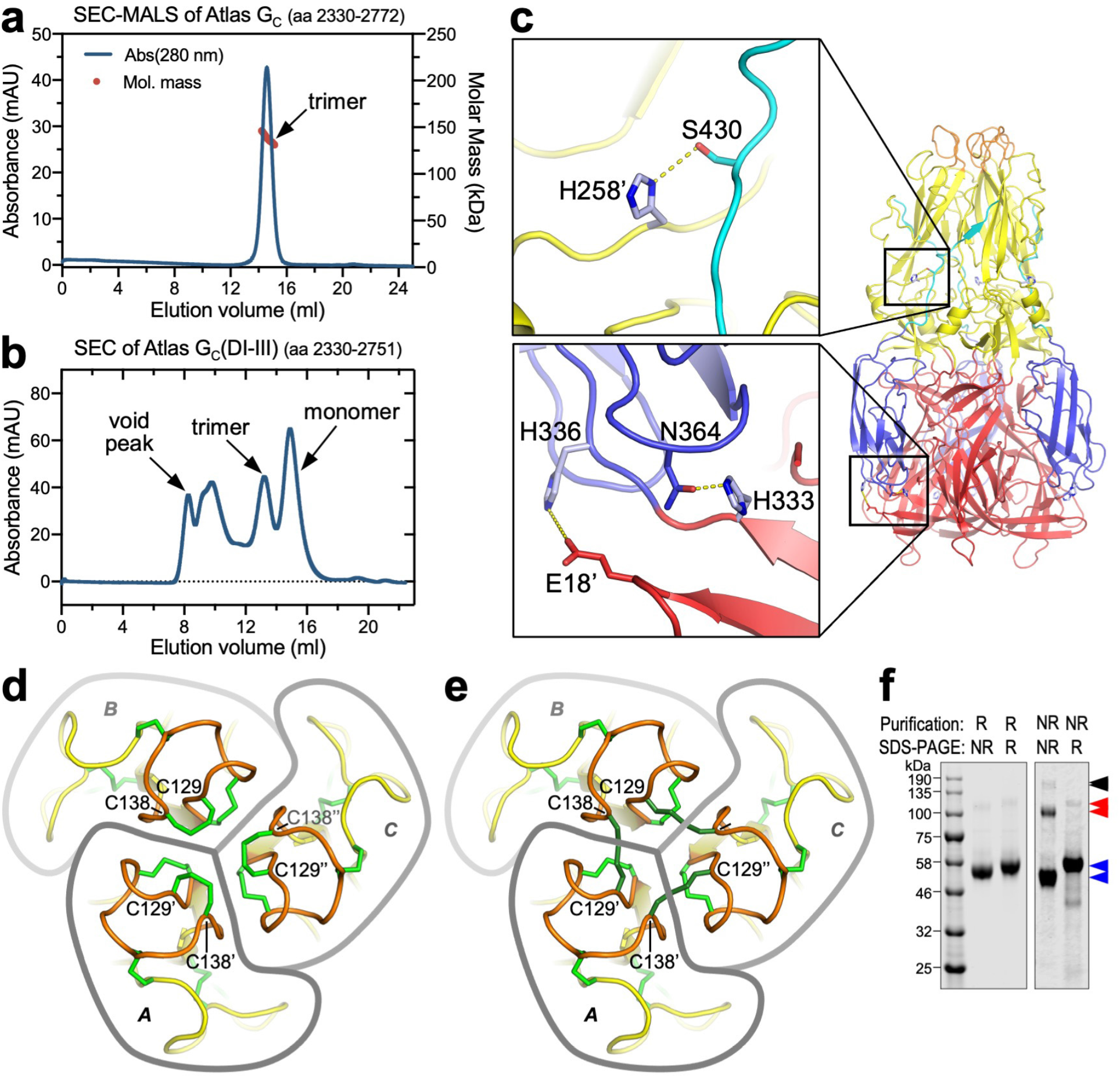
Oligomeric states and disulfide bonding pattern of Atlas G_C_. **(a)** SEC-MALS of Atlas G_C_ ectodomain (residues 2330-2772). The protein formed stable trimers. **(b)** SEC of Atlas G_C_(DI-III) (residues 2330-2751) is expressed in a mixture of oligomeric states, including monomers, trimers and larger aggregates (void peak). G_C_(DI-III) monomers were unstable. **(c)** His258, His333 and His336 form interprotomer or interdomain polar contacts predicted to stabilize the G_C_ trimer specifically at pH <6, when the histidine side chains are charged. Residues from different protomers are denoted with a prime symbol. **(d-e)** Views along the threefold axis of the G_C_ trimer without, (**d**), and with, (**e**), intermolecular disulfides modeled between the cysteines most proximal to the axis, Cys129 and Cys138. **(f)** Coomassie-stained SDS-PAGE gels of Atlas G_C_ purified under non-reducing conditions (NR) or with 0.5 mM TCEP reducing agent (R). Gel samples were prepared with either non-reducing (NR) or reducing (R) SDS-PAGE loading buffer. Approximately 15% of the Atlas G_C_ sample purified under non-reducing conditions contained intermolecular disulfides. Arrowheads indicate bands corresponding to monomers (blue), dimers (red) and trimers (black).

### pH-dependent stabilization of the Atlas G_C_ trimer by protonated histidine residues

The increase in positive charge resulting from histidine protonation is an important part of the pH sensing mechanism of viral class II fusion proteins. Protonation of conserved histidines at the domain I-domain III interface of alphavirus, flavivirus and phlebovirus glycoproteins promotes the fusogenic conformational change by destabilizing the prefusion conformation and stabilizing the postfusion conformation (*19, 27, 58-61*). For example, histidines in domain III of phlebovirus G_C_ proteins form interprotomer salt bridges with negatively charged side chains in the postfusion trimer and mutation of one such histidine in RVFV, His1087, renders the virus uninfectious (*62*). Similarly, in Atlas G_C_ His258, His333 and His336 form interprotomer or interdomain polar contacts (**Fig. 5c**). By stabilizing the trimeric postfusion-like conformation of Atlas G_C_ specifically at low pH these histidine-dependent contacts would reduce the reversibility of any preceding conformational change in the protein in acidic endosomal compartments. The striking parallels in how Atlas G_C_ and class II fusogens from infectious viruses respond at the ultrastructural level to environmental cues support the hypothesis that Atlas G_C_ would have membrane fusion activity in late endosomes, like phleboviruses and many retroviruses.

### Fifteen disulfide bonds stabilize Atlas G_C_ in its postfusion-like conformation

With 30 cysteine residues forming 15 disulfide bonds, Atlas G_C_ contains twice the average abundance of cysteines, more than has been found in any other class II protein. 20 of these cysteines form disulfides that are structurally conserved in phleboviruses (including RVFV, DABV and HRTV) but not in other class II proteins (**Supplementary Fig. 5**). An eleventh disulfide, in domain III, is conserved in Atlas virus, DABV and HRTV but not RVFV. However, Atlas G_C_ contains four additional disulfides not conserved in phleboviruses: one in the fusion loop, two in domain II in the *β*-hairpin containing the *ij* loop (one of the cysteines forming these disulfides is conserved in phleboviruses but forms a disulfide with a cysteine in a different *β*-strand in domain II), and one in domain III (**Figs. 2c, 3a; Supplementary Fig. 5**). The Atlas-specific disulfide in the fusion loop, Cys129-Cys138, appears likely to confer novel biochemical properties. As discussed above, the side chains of Cys129 and Cys138 extend the hydrophobic surface formed by conserved residues in the fusion loop that are required for membrane insertion. Furthermore, due to the location of Cys129 and Cys138 close to the threefold symmetry axis of the trimer, the side chains of the two residues can be rearranged by torsional rotation, without changes to the polypeptide backbone, to form intermolecular disulfides across the trimer interface, thereby crosslinking all three protomers in the trimer (**Fig. 5d,e**). SDS-PAGE under non-reducing conditions showed that approximately 15% of protein purified without reducing agents contained one or more intermolecular disulfide bonds. However, under the reducing conditions used to purify Atlas G_C_ for cryo-EM imaging (0.5 mM TCEP), no crosslinked protein was detected (**Fig. 5f**). Exchange of the Cys129-Cys138 bonds from intramolecular to intermolecular disulfides could provide a novel mechanism to stabilize the Atlas G_C_ trimer in its postfusion-like conformation and reduce the reversibility of any preceding conformational change. Disulfide-mediated crosslinking of the postfusion conformation has not been described in any viral or non-viral membrane fusion proteins but could potentially function as a mechanism to promote a fusogenic conformational change in Atlas G_C_ or in infectious viruses with fusion proteins related to Atlas G_C_.

### Atlas G_C_ retains membrane fusion activity

Having identified the hallmarks of a fusion protein in Atlas G_C_, we measured its membrane fusion activity in a cell-cell fusion assay. CHO cells were transfected with plasmids encoding Atlas G_C_ or Atlas G_N_-G_C_. To promote plasma membrane localization and minimize ER retention, we replaced the predicted transmembrane domain and cytosolic tail of G_C_ with the C-terminal transmembrane anchor and cytosolic tail from HLA-A2, known to localize to the plasma membrane (*63*). Plasmids encoding vesicular stomatitis virus (VSV) G, or no protein, were used as positive and negative controls, respectively. Since Atlas G_C_ trimers require low pH and BMP or PS to efficiently bind membranes, we treated transfected cells with exogenous BMP and then transferred them to pH 4.5 buffered medium to trigger fusion. Similar treatments have been used previously to measure cell-cell fusion activity of flavi- and alphaviruses (*46*). Confocal light microscopy with nuclear and plasma membrane stains showed that cells with three or more nuclei were common in cells expressing Atlas G_C_ following treatment with BMP and pH 4.5, though less abundant than in cells expressing VSV G, with or without treatment (**Fig. 6a**). Cells with more than three nuclei were rare in cells expressing Atlas G_N_-G_C_. To quantify cell-cell fusion we counted nuclei and multinuclear cells in micrographs. We found that in cells expressing Atlas G_C_, the fraction of multinuclear cells, defined as cells containing two or more nuclei, was 25 ± 3 % following treatment with BMP and pH 4.5, versus 10 ± 0.2 % without treatment (**Fig. 6b**). Among cells expressing Atlas G_N_-G_C_, 13 ± 1 % were multinuclear with treatment, versus 10 ± 1 % without treatment. By comparison, in cells expressing VSV G, 73 ± 8 % of treated cells and 41 ± 8 % of untreated cells were multinuclear. 8 ± 1 % of cells transfected with empty vector were binuclear with or without treatment, but none contained more than two nuclei (**Supplementary Fig. 9**). We conclude that Atlas G_C_ has approximately one third of the membrane fusion activity of VSV G under the treatment conditions tested, which is significant given that VSV G is considered highly fusogenic and widely as a model fusogen. The Atlas G_N_-G_C_ construct tested had little fusion activity. The reason for this remains unclear – possible explanations include incomplete proteolytic processing at the G_N_-G_C_ junction, lower expression level, or absence of a priming factor in CHO cells.

**Figure 6.**
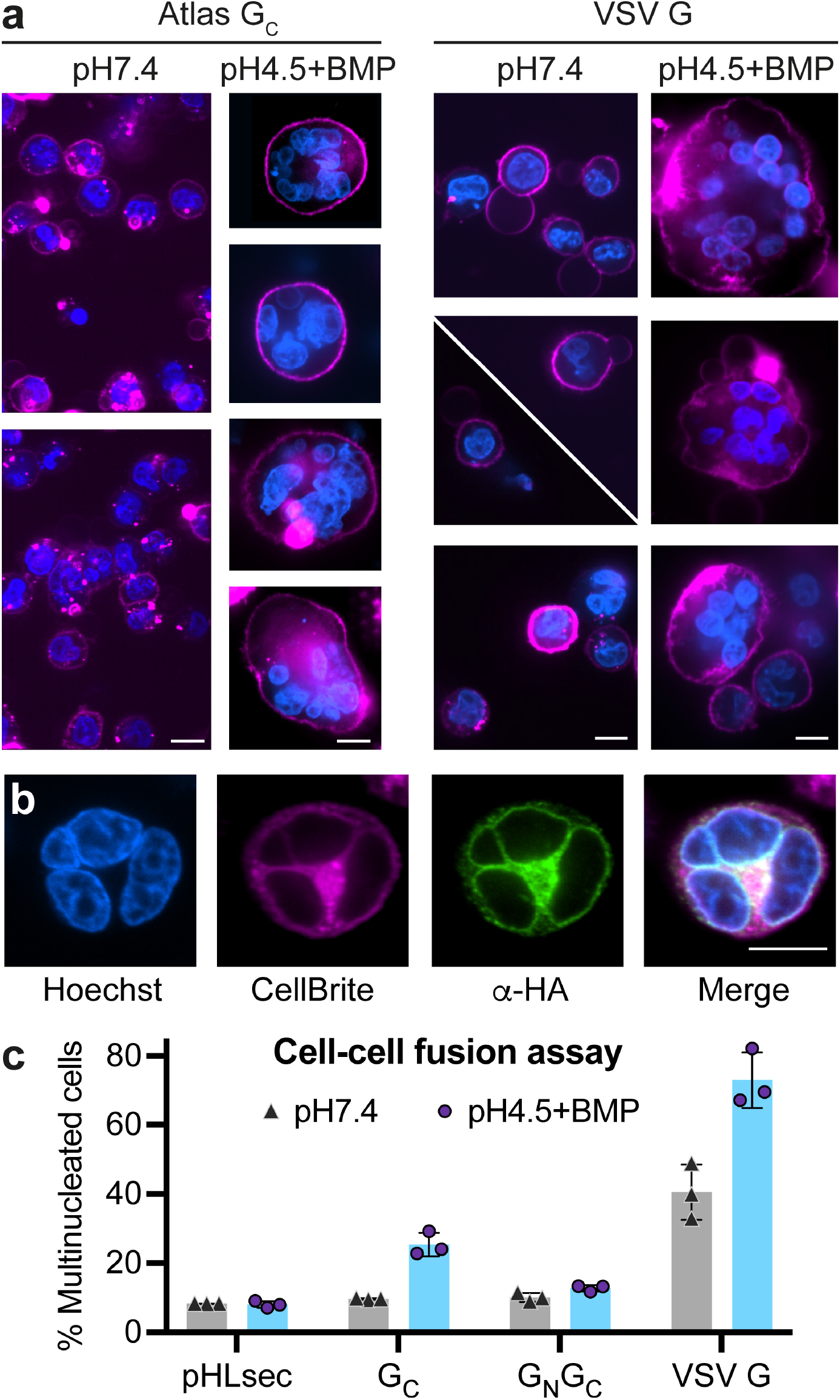
Cell-cell fusion assay with cells transfected with Atlas G_C_ or G_N_-G_C_. **(a)** Confocal micrographs of CHO cells expressing Atlas G_C_ fused to the C-terminal transmembrane anchor from HLA-A2, or VSV G. Cells were treated with pH 4.5 buffer containing BMP, or pH 7.4 containing no exogenous lipid. Blue, Hoechst 33342 nuclear stain; Magenta, CellBrite Red plasma membrane dye. Micrographs were selected to show fusion events – see **Supplementary Fig. 9** for representative raw micrographs. **(b)** Confocal micrograph of a multinucleated CHO cell expressing Atlas G_C_ following treatment with pH 4.5 and BMP. Blue, Hoechst 33342; magenta, CellBrite Red; green, *α*-HA antibody for Atlas G_C_ detection. All scale bars in (**a**) and (**b**) are 10 µm. **(c)** The fraction of multinuclear cells transfected with plasmids encoding Atlas G_C_, Atlas G_N_-G_C_, VSV G or no protein (pHLsec empty vector control) was calculated by counting mono- and multinucleated cells in confocal micrographs, using the Hoechst and CellBrite stains. Error bars represent standard deviation between measurements, n = 3. See **Supplementary Dataset 2** for source data.

### Transcriptional activity of Atlas virus

The transcription of endogenous retroviruses is tightly regulated to prevent harmful gene expression and genome damage from transposition events. However, some ERVs escape repression and are highly transcribed, including a subset of belpaoviruses from the parasitic trematode *Schistosoma mansoni* (blood fluke) (*64*). As a first step to assess the expression level of Atlas virus in *A. ceylanicum*, we analyzed transcript levels of Atlas and other intact belpaoviruses in previously published transcriptional profiling (RNA-Seq) data (*65*). Transcript abundance of the nine belpaovirus ERVs encoding complete and intact polyproteins (**Fig. 1a**) varied, ranging from undetectable to above the median value for all annotated genes (**Fig. 7, Supplementary Fig. 10**). For Atlas virus, transcript levels varied across developmental stages: expression was above the median for annotated genes at the larval L3 stage, below the first quartile at stage L4, and slightly below the median in adult hookworms (**Fig. 7a**). More precisely, Atlas virus transcript expression increased 15-fold from the L4 to adult (mixed sex) stages (adjusted *p*-value <0.05, see **Supplementary Dataset 3** for differential expression calculations). Because the L3 sample was not replicated, statistical significance could not be established for the differences in expression between L3 and other stages. RNA-Seq reads coverage over the Atlas virus gene was greater in mixed sex adults than in L4 larvae (**Fig. 7b**). In contrast, the difference in Atlas transcription from L4 to male adults was not statistically significant. This suggests the increase in expression from L4 to mixed sex adults was primarily driven by increased expression in female tissues or unlaid eggs. In conclusion, Atlas virus is transcriptionally active at specific stages of hookworm development.

**Figure 7.**
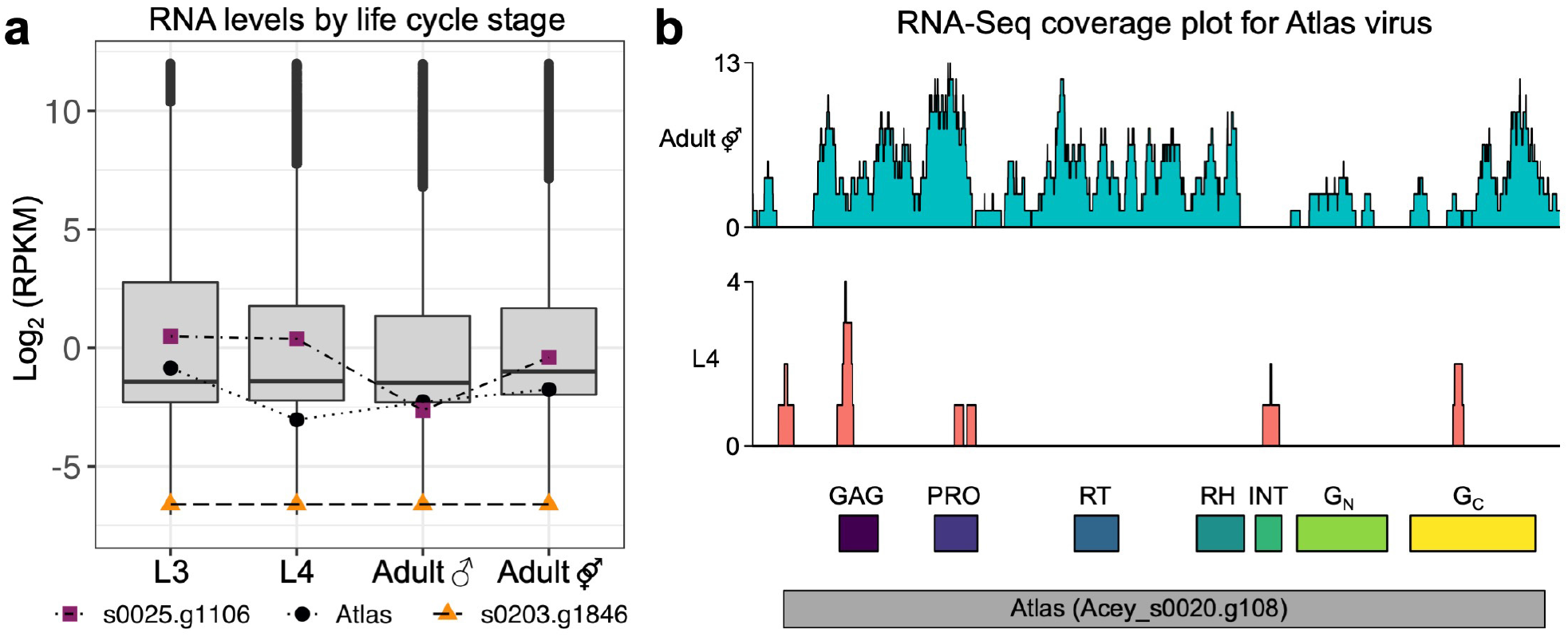
Transcriptional activity of Atlas virus in *A. ceylanicum* hookworm. **(a)** Boxplot representation of RNA-Seq expression in Reads Per Kilobase of transcript per Million mapped reads (RPKM) for all annotated genes. The RPKM values for Atlas virus and two other representative intact *A. ceylanicum* belpaovirus ERVs at different developmental stages (L3 and L4 larva, adult male and adult mixed sex) are highlighted. RNA-Seq data are from Bernot *et al.* (*65*). Trendlines were added to aid interpretation. **(b)** Distribution of raw RNA-Seq reads over the Atlas virus (*Acey_s0020.g108*) coding sequence for the Adult (mixed sex) and L4 developmental stages. A snapshot of the same region from the Integrative Genomics Viewer (IGV) with individual unmerged BAM files is shown in **Supplementary Fig. 10**.

## Discussion

Here, we identify Atlas virus as an intact ERV with novel features in the human hookworm *A. ceylanicum*. The cryo-EM structure of the Atlas Env reveals a class II viral fusion protein fold remarkably similar to that of the G_C_ glycoprotein from Rift Valley Fever virus. Atlas G_C_ has the hallmarks of an active class II membrane fusion protein: a stable trimeric assembly, a putative fusion loop, membrane insertion triggered by low pH with specificity for late endosomal lipid composition, and membrane fusion activity. Moreover, Atlas virus mRNA is expressed at specific stages of hookworm development.

The envelope proteins from retroviruses, including ERVs, that have been biochemically characterized were all found to be class I fusion proteins with an *α*-helical coiled-coil as the core fold. Viral class II fusion proteins have so far been found only in non-integrating RNA viruses. Our discovery of an ERV with a functional, phlebovirus-like class II fusion protein that is structurally unrelated to other retrovirus Envs reveals an unexpected degree of structural and genetic plasticity in retroviruses. Our work supports the model first proposed based on phylogenetic studies that the nematode belpaoviruses acquired their Env by horizontal gene transfer from a virus from the *Phenuiviridae* family or a phlebovirus-like ancestor(*13*). It remains unclear whether RNA encoding the phlebovirus-like glycoprotein integrated into the belpaovirus ancestor as mRNA in a splicing event, or by first becoming a substrate for the reverse transcriptase with subsequent genomic integration as dsDNA.

While rare, horizontal gene transfer of atypical fusogens into retroelements is not unique to the belpaoviruses. The Tas element from the nematode *Ascaris lumbricoides*, the most common parasitic worm in humans, has an *env* gene with weak genetic similarity to herpesvirus gB proteins (*13*), which have a class III fusion protein fold (also found in *Rhabdoviridae* and *Baculoviridae*). Taken together, these findings lead us to hypothesize that acquisition of a fusion protein from an infectious virus represents a general paradigm for how retrotransposons can become retroviruses, and indeed how ancestral reverse-transcribing viruses may have originated.

The *env* gene is often first element to be lost in ERVs, as it is not required for intracellular proliferation, so it is notable that the G_N_-G_C_ *env* module is intact in Atlas virus. With its identical LTRs, and no stop codons or frameshift mutations, Atlas virus shows all the signs of being intact and recently active. This supports the notion that the envelope may be functional (*11*). The preserved biological activities of Atlas G_C_ suggest these activities could have cellular functions in health and disease, as reported for a small but increasing number of other ERV *env* and *gag* gene products (*1, 3–5*). Further studies are required to determine the full extent to which protein expression from transposable elements – and its dysregulation – contribute to basic cellular functions, embryonic development and disease outcomes. This work provides a blueprint for such efforts.

## Materials and Methods

### Sequence analysis of *A. ceylanicum* Atlas virus

A PSI-BLAST search (*33*) for protein sequences similar to biochemically characterized phlebovirus fusion proteins identified the gene *Acey_s0020.g108* (UniProt A0A016UZK2; Genbank JARK01001356.1, genomic translation EYC20859.1) in the human hookworm *Ancylostoma ceylanicum* as the most similar sequence outside infectious virus taxa, with an Expected value (E-value) of 10^-20^ against the RVFV G_C_ sequence. A second iteration performed using position-specific scoring matrix based on an alignment of sequences identified in the first iteration gave an E-value of 10^-144^.

### Protein expression and purification

Synthetic genes encoding soluble ectodomain fragments of the Env of endogenous retrovirus Y032_0020g108 from *A. ceylanicum* were subcloned into the pMT/BiP/V5-His vector (ThermoFisher) in frame with the BiP signal sequence and the C-terminal V5 and six-histidine tags. The constructs referred to here as Atlas G_C_ and Atlas G_C_(DI-III) span amino acids 2330-2772 and 2330-2751 from UniProt A0A016UZK2, respectively. Atlas G_C_ mutants were generated by Dpn I-based site-directed mutagenesis. D.mel-2 insect cells (ThermoFisher) were co-transfected with the expression construct and blasticidin resistance marker pCoBlast (ThermoFisher) at a 20:1 molar ration and cultured for 6 weeks in 0.5 µg ml^-1^ blasticidin to obtain a population of expressor cells. Expression was induced in a shaking cell suspension at 27°C with 0.5 mM CuSO_4_ at a cell density of 5 x 10^6^ cells ml^-1^. The cell culture medium was harvested 4 to 5 days after induction, centrifuged to remove cells (2,000 g) and cell debris (17,000 g), filtered with a 0.2 µm filter, concentrated by tangential-flow filtration, and buffer exchanged into 20 mM Tris pH 7.8, 0.3 M NaCl, 5% glycerol, 20 mM imidazole, 0.5 mM TCEP (tris(2-carboxyethyl)phosphine). Atlas G_C_ was purified by nickel-affinity chromatography with a HisTrap Excel column (Cytiva), followed by anion-exchange chromatography with a MonoQ or Resource Q column (Cytiva) using 20 mM Tris pH 8.0, 50 mM NaCl, 5% glycerol, 0.5 mM TCEP as the binding buffer and binding buffer plus 1 M NaCl as the elution buffer. Peak fractions were concentrated and further purified by size-exclusion chromatography (SEC) with a Superdex 200 Increase (10/300) column (Cytiva) in 20 mM Tris pH 7.8-8.0, 0.15 M NaCl, 5% glycerol and 0.5 mM TCEP. The C-terminal V5 and histidine tags were optionally cleaved by incubation with carboxypeptidase A (CPA) for 3 h at 4°C (1:500 CPA:G_C_ molar ratio).

### Liposome binding assay

1-palmitoyl-2-oleoyl-sn-glycero-3-phosphocholine (PC), 1-palmitoyl-2-oleoyl-snglycero-3-phosphoethanolamine (PE), egg sphingomyelin (SM), 1-palmitoyl-2-oleoyl-sn-glycero-3-phospho-L-serine (PS), 1-palmitoyl-2-oleoyl-sn-glycero-3-phospho-(1’-rac-glycerol) (PG), (*S*,*R*) Bis(Monoacylglycero)Phosphate (BMP) (Avanti Polar Lipids), and 1-cholesterol (Sigma-Aldrich) were dissolved in chloroform. 25 mM lipid solutions were mixed at various molar ratios and dried under nitrogen gas for over 4 h. The lipid film was resuspended in Liposome Buffer (20 mM Tris pH 7.8, 0.15 M NaCl, 5% glycerol, 0.5 mM TCEP, 2 mM MgCl_2_, 2 mM CaCl_2_, 2 mM KCl) and subjected to five cycles of freeze-thawing in liquid nitrogen, followed by 25 cycles of extrusion through two 0.2 µm polycarbonate filter membranes (Whatman). Purified Atlas G_C_ was added in a 1:771 protein:lipid molar ratio and incubated at 37°C for 5 min. The pH was reduced by adding a 2 M stock solution of sodium acetate pH 4.6 or 4.0 to a final concentration of 0.2 M. Following a 2 h incubation at 37°C the pH of the suspension was neutralized with 1 M Tris pH 8. OptiPrep density gradient medium (Sigma-Aldrich) was added to a concentration of 40%, maintaining 0.15 M NaCl throughout. Approximately 0.5 ml of the liposome suspension was placed in a centrifuge tube, overlayed with a 2.5 ml cushion of 30% OptiPrep solution, and centrifuged at 100,000 g for 1 h at 4°C in a TLA100.3 rotor (Beckman Coulter). Top (T) and bottom (B) fractions (approximately 1.5 ml each) were collected from the top meniscus with a micropipette. Atlas G_C_ was quantified by densitometry of the absorbance at 700 nm of bands in Coomassie-stained SDS-PAGE gels with an Odyssey scanner (LI-COR). Flotation was defined as the amount of Atlas G_C_ in the top fraction divided by the total amount of Atlas G_C_ in both fractions.

For measurement of liposome diameter by dynamic light scattering (DLS), the liposome suspensions were diluted ten-fold in Liposome Buffer prior to the addition of 40% OptiPrep solution. Following centrifugation liposome diameters were measured in 384-well clear-bottomed optical imaging plates (Corning) with a DynaPro Plate Reader III (Wyatt Technologies). The mean diameter was calculated as the average of three independent measurements, each consisting of fifteen two-second acquisitions. Protein-free acidified liposome controls were treated and measured in parallel, with Liposome Buffer instead of Atlas G_C_ solution.

### Size-exclusion chromatography and multi-angle scattering (SEC-MALS) analysis

100 µl samples containing 1.6 – 2.5 mg ml^-1^ Atlas G_C_ were analysed by size-exclusion chromatography (SEC) at 293 K on a Superdex 200 (10/300) column (Cytiva) in 20 mM Tris pH 7.8, 0.15 M NaCl, 5% glycerol and 0.5 mM TCEP with a flow rate of 0.5 ml min^-1^. The SEC system was coupled to both multi-angle light scattering (MALS) and quasi-elastic light scattering (QELS) modules (DAWN-8+, Wyatt Technology). The protein was also detected as it eluted from the column with a differential refractometer (Optilab T-rEX, Wyatt Technology) and a UV detector at 280 nm (Agilent 1260 UV, Agilent Technology). Molar masses of peaks in the elution profile were calculated from the light scattering and protein concentration, quantified using the differential refractive index of the peak assuming a dn/dc of 0.1860, with ASTRA6 (Wyatt Technology).

### Cryo-EM sample preparation and data collection

Purified Atlas G_C_ trimer (3 μl at a concentration of 0.025 mg ml^-1^) in 20 mM Tris pH 7.8, 0.15 M NaCl, 5% glycerol and 0.5 mM TCEP, was applied onto glow-discharged R1.2/1.3 400 mesh copper grids (Quantifoil Micro Tools, Germany). The grids were blotted for 4 s and plunge-frozen in liquid ethane with a Vitrobot Mark IV (ThermoFisher) at 4°C and 100% humidity. Preliminary sample screening and initial datasets were acquired on a FEI Tecnai F20 microscope operated at 200 kV equipped with Falcon II direct electron detector (ThermoFisher) at -4 µm defocus. High-resolution cryo-EM dataset collection was performed on a Titan Krios microscope (ThermoFisher) operated at 300 kV equipped with a 20 eV slit-width GIF quantum energy-filtered Gatan K2-Summit direct electron detector in counting mode. A total of 3,027 movies were recorded at a calibrated magnification of 130,000x leading to a magnified pixel size of 1.047 Å on the specimen. Each movie comprised 36 frames with an exposure rate of 1.28 e^-^Å^-2^ per frame, with a total exposure time of 8 s and an accumulated exposure of 46.18 e^-^ Å^-2^. Data acquisition was performed with EPU Automated Data Acquisition Software for Single Particle Analysis (ThermoFisher) with three shots per hole at -1.3 µm to -3.5 µm defocus.

### Image processing

Micrographs from initial datasets allowed us to obtain a consistent model at ∼19 Å resolution from 3,790 particles selected after 2D and 3D classification, and consequent auto-refinement. All movies from high-resolution datasets were motion-corrected and dose-weighted with MOTIONCOR2 (*66*). Aligned, non-dose-weighted micrographs were then used to estimate the contrast transfer function (CTF) with GCTF (*67*). All subsequent image processing steps were performed using RELION 3.0 (*68, 69*). 2D references from initial datasets were used to auto-pick the micrographs. One round of reference-free 2D classification was performed to produce templates for better reference-dependent auto-picking, resulting in a total of 987,570 particles. After a first round of 2D classification, 595,011 particles were selected to perform a second 2D classification, resulting in a final number of 320,041 selected particles. Then, a 3D classification imposing C3 symmetry was performed using the model from the initial datasets filtered at 40 Å resolution as initial model. The best class, containing 197,145 particles, was selected and subjected to 3D auto-refinement imposing C3 symmetry, yielding a map with an overall resolution at 4.11 Å based on the gold-standard (FSC = 0.143) criterion. After refinement, the CTF refinement (per-particle defocus fitting and beam tilt estimation) and Bayesian polishing (*70*) routines implemented in RELION 3.0 were performed, yielding a final map with an overall resolution at 3.76 Å. Local resolution was estimated with RELION.

### Model building and refinement

The most similar sequence to Atlas G_C_ with a structure available was glycoprotein G_C_ from Rift Valley Fever virus (RVFV). The crystal structure of RVFV G_C_ in the postfusion conformation (PDB:6EGU (*27*)) was used as template to build a homology model with the sequence of Atlas G_C_ using the Swiss-Model server (https://swissmodel.expasy.org). The output model was docked as a rigid body into the density with UCSF Chimera (*71*). Initial docking was performed manually and was followed by real space fitting with the Fit in Map routine. A preliminary step of real space refinement was performed on the three-subunit model, with Phenix 1.13 (*72*), with global minimization, atomic displacement parameter (ADP), simulated annealing and morphing options selected. The model was then rebuilt in Coot (*73*) to optimize the fit to the density. Due to low resolution information in the fusion loop region, the density was converted to .mtz file using CCP-EM software package tools and blurring of the density allowed us to localize bulky residues and disulphide bonds, and thus use them as a guide to build the entire fusion loop. A final step of real space refinement was performed with Phenix 1.15, with global minimization and ADP options selected. The following restraints were used in the real space refinement steps: secondary structure restraints, non-crystallographic symmetry (NCS) restraints between the protein subunits, side chain rotamer restraints, and Ramachandran restraints. Key refinement statistics are listed in **Supplementary Table 1**.

### Model validation and analysis

The FSC curve between the final model and full map after post-processing in RELION, Model vs Map, is shown in **Supplementary Fig. 3**. Cross-validation against overfitting was performed as described (*74*). The atoms in the final atomic model were displaced by 0.5 Å in random directions with Phenix. The shifted coordinates were then refined against one of the half-maps generated in RELION, the “work set”. This test refinement was performed in Phenix using the same procedure as for the refinement of the final model (see above). The other half-map, the “test set” was not used in refinement for cross-validation. FSC curves of the refined shifted model against the work set, FSCwork, and against the test set, FSCtest, are shown in **Supplementary Fig. 3**. The FSCwork and FSCtest curves are not significantly different, consistent with the absence of overfitting in our final models.

The quality of the atomic models, including basic protein geometry, Ramachandran plots, clash analysis, was assessed and validated with Coot, MolProbity (*75*) as implemented in Phenix 1.15, and with the Worldwide PDB (wwPDB) OneDep System (https://deposit-pdbe.wwpdb.org/deposition).

### Cell-cell fusion assay

Chinese hamster ovary (CHO) Lec3.2.8.1 cells were transfected with pHL-Sec plasmids encoding ectodomain fragments of Atlas G_N_-G_C_ (polyprotein residues 1909-2795) or Atlas G_C_ (residues 2330-2795) fused to the C-terminal transmembrane domain from HLA-A2 (residues 288-345) and cloned in frame with the vector’s secretion signal and a C-terminal HA tag. Empty pHL-Sec plasmid, and pcDNA encoding vesicular stomatitis virus (VSV) G, were used as negative and positive controls, respectively. 16-20 h post-transfection, cells were transferred to PBS supplemented with 2.5 mM BMP (18:1(*S,S*) Bis(Monoacylglycero)Phosphate (BMP); Avanti Polar Lipids). To obtain a homogeneous BMP suspension, the mixture was freeze-thawed five times using liquid nitrogen and a water bath, followed by a 3 min incubation in a sonicating water bath. Cells were incubated in the BMP suspension (or PBS for the untreated control) at 37°C for 5 min, shown previously to be sufficient for anionic lipid incorporation into the plasma membrane (*46*). Cell were transferred to pH 4.5 complete medium (DMEM adjusted to pH 4.5 with HCl supplemented with 10% foetal bovine serum), or pH 7.4 complete medium for the untreated control, and centrifuged at 2,500 g at 37°C for 2 min. Cells were immediately resuspended in complete media and plated out. Following reattachment (4-6 h after treatment), cells were stained with Hoechst 33342 (Bio-Rad) and CellBrite Red cytoplasmic membrane dye (Biotium, cat. no. 30023), and imaged on a Nikon iSIM Swept Field inverted confocal microscope with a 60x/1.2NA water objective. Cluster analysis of the Hoechst channel with Fiji (*76*) was used to count single nuclei and identify polynuclear clusters. Nuclei within polynuclear clusters were counted by visual inspection. The plasma membrane stain was used to confirm polynuclear clusters and count the number of multinuclear cells, defined as cells with two or more nuclei, by visual inspection. The fraction of multinucleated cells (*F*) was calculated using the formula:

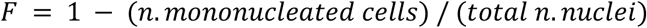

### RNA-Seq analysis

We analysed data published by Hawdon and colleagues (*65*). ArrayExpress run accession numbers and corresponding life cycle stages were: SRR6359160, L3; SRR6359161 and SRR6359163, L4; SRR6359164 and SRR6359165, Adult mixed pooled worms; SRR6359162 and SRR6359166, Adult male pooled worms. Our mapping and counting strategy was modified as recommended for repetitive genomic features (*77*). Reads were mapped against the reference *A. ceylanicum* genome (*78*) with STAR v2.7.5a (*79*) with parameters: “--outSAMtype BAM SortedByCoordinate --runMode alignReads --outFilterMultimapNmax 1000 --outSAMmultNmax 1 --outFilterMismatchNmax 3 --outMultimapperOrder Random --winAnchorMultimapNmax 1000 --alignEndsType EndToEnd --alignIntronMax 1 --alignMatesGapMax 350”. FeatureCounts v2.0.1 (*80*) with parameters “-M -F GFF -s 0 -p -t exon -g gene_id” was used to count reads over a modified annotation file based on the original annotation (*78*) and downloaded from Wormbase Parasite (*81*). The original annotation file in GFF format was slightly modified to contain a “gene_id” feature in the ninth column and hence facilitate the calculation of aggregate reads by gene (for more details see header of the GFF file, **Supplementary Dataset 4**). The resulting table of counts was imported into R (*82*) and processed for differential expression analysis using DESeq2 v1.26.0 (*83*). Plots were generated using ggplot2 (*84*) and karyoploteR (*85*). All scripts and instructions for how to use them for RNA-Seq analysis are available from Github [https://github.com/annaprotasio/Merchant_et_al_2020].

### Statistics

Error bars represent the standard deviation or standard error –as indicated in the respective figure legend– of 2-7 replicates conducted across at least two independent experiments. SDS-PAGE gels and DLS data shown are representative of at least two independent experiments. Significance was determined by 2-way ANOVA analysis using Sidak’s multiple comparisons test with a 95% confidence interval, in Prism 8 (GraphPad). **, 0.001 < p < 0.01; *, p < 0.05; ns, not significant. Source data are provided in **Supplementary Datasets 1 and 2.** No statistical methods were used to predetermine sample size, experiments were not randomized, and the investigators were not blinded to experimental outcomes.

### Data availability

The atomic coordinates were deposited in the Protein Data Bank with code 7A4A [https://doi.org/10.2210/pdb7A4A/pdb]. The cryo-EM density was deposited in the EM Data Bank with code EMD-11630 [https://www.ebi.ac.uk/pdbe/entry/emdb/EMD-11630]. The raw electron micrographs were deposited in EMPIAR with code EMPIAR-10266 [https://dx.doi.org/10.6019/EMPIAR-10266]. RNA-Seq analysis scripts and instruction are available from Github [https://github.com/annaprotasio/Merchant_et_al_2020]. Other data are available from the corresponding author upon reasonable request.

## Acknowledgements

We acknowledge the MRC Laboratory of Molecular Biology Electron Microscopy (EM) and UK national electron Bio-Imaging Centre (eBIC) at Diamond Light Source (DLS) for access and support of EM sample preparation and data collection. We acknowledge DLS for access and support of eBIC under proposals EM17434 and EM20121 funded by the Wellcome Trust, MRC and BBSRC. We are grateful to Kyle Dent for providing assistance in using the microscopes at eBIC. We are grateful for access to the MRC-LMB Scientific Computing facilities. This work was supported by Senior Research Fellowships 101908/Z/13/Z and 217191/Z/19/Z from the Wellcome Trust and NIH grant R01 GM102869-01 to Y.M.

## Author contributions

M.M. and Y.M. conceived the experiments. M.M. purified the proteins and performed the biochemical assays. C.P.M collected and processed the cryo-EM data, and performed the image reconstruction. C.P.M and Y.M. built and refined the atomic model. Y.L. and H.Z. carried out the cell-cell fusion assay. A.V.P performed the RNA-Seq analysis. All authors contributed to the figures. Y.M. wrote the manuscript. Project supervision, administration and funding acquisition, Y.M.

## Materials & Correspondence

Correspondence and material requests should be addressed to Y.M.

## Supplementary Information

**Supplementary Fig. 1.**
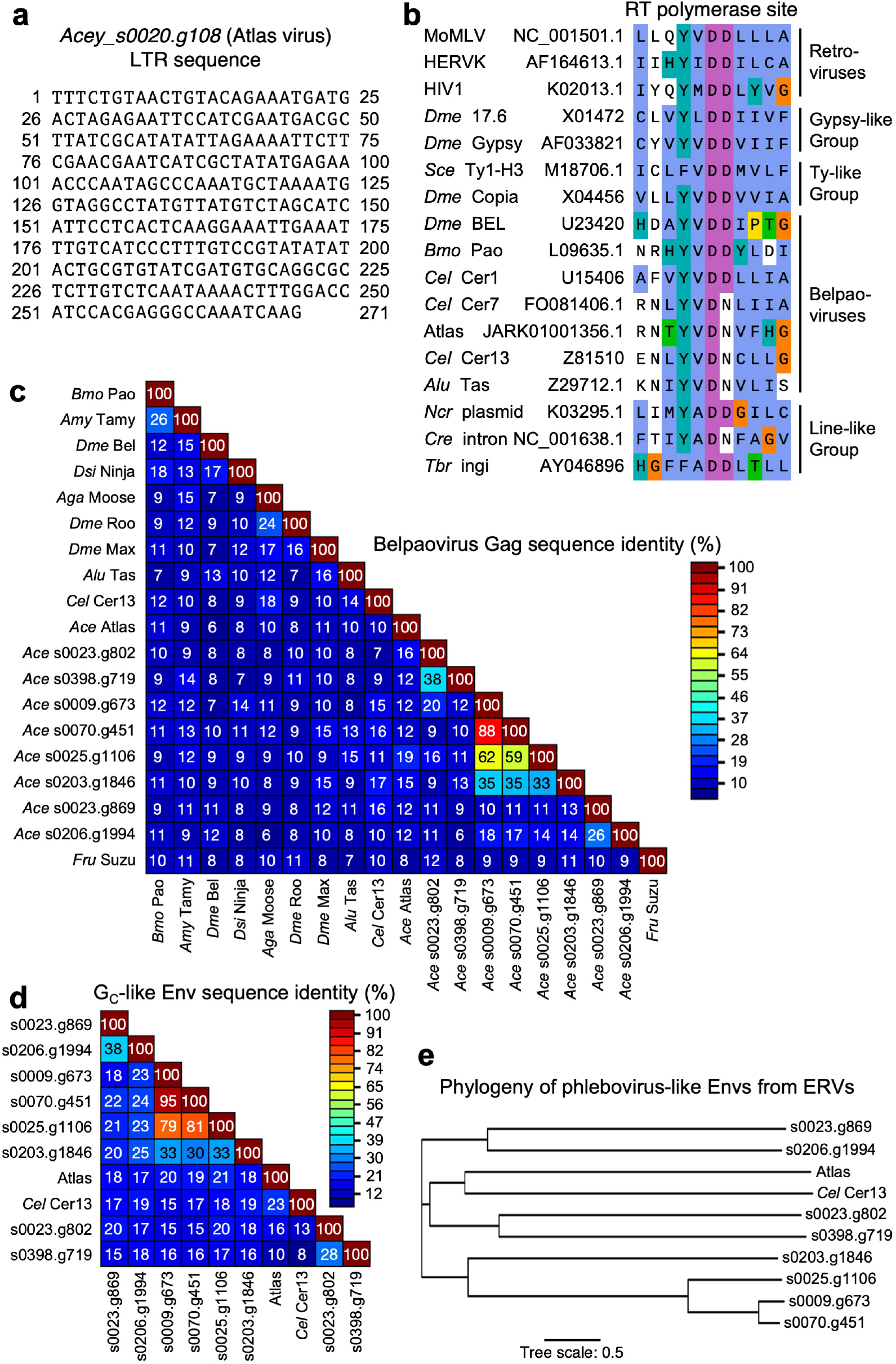
Atlas is an intact belpaovirus with a phlebovirus-like Env in the hookworm *A. ceylanicum*. **(a)** Nucleotide sequence of the Atlas virus LTRs, predicted by REPuter(*86*). The sequences of the two LTRs is 100% identical. **(b)** Aligned protein sequences of reverse transcriptase (RT) polymerase site from Atlas and other representative belpaoviruses and retroviruses. Endogenous belpaoviruses in *A. ceylanicum* have an aspartate to asparagine substitution (Y[X]DD to YVDN) in the most conserved reverse transcriptase motif, the polymerase site (Motif V or Motif C). Nucleotide accession numbers are listed after virus names. MoMLV, Murine Moloney leukemia; HERVK, human ERV-K; *Dme*, *D. melanogaster*; *Sce, Saccharomyces cerevisiae*; *Bmo, Bombyx mori*; *Cel, C. elegans*; *Ace, A. ceylanicum*; *Alu, Ascaris lumbricoides*; *Ncr, Neurospora crassa*; *Cre, C. reinhardtii*; *Tbr, Trypanosoma brucei*. **(c)** Matrix of sequence identities for pairs of Gag protein sequences from representative viruses from the *Belpaoviridae* family and the belpaovirus ERVs from *A. ceylanicum* with intact protein coding regions. *Amy, Antheraea mylitta; Dsi, Drosophila simulans; Aga, Anopheles gambiae; Fru, Fugu rubripes.* With less than 20% Gag protein sequence identity to its closest homolog (*87*) (*Acey_s0020.g1106*), Atlas virus is a candidate for classification as new member of the *Belpaoviridae* family (which contains a single genus, *Semotivirus*). **(d)** Sequence identity matrix for phlebovirus-like Env protein sequences from nematode belpaoviruses. Identity matrices were calculated with SDT (*88*). **(e)** Phylogenetic tree of phlebovirus-like Env proteins from nematode belpaoviruses. Drawn with Interactive Tree Of Life (iTOL) (*36*).

**Supplementary Fig. 2.**
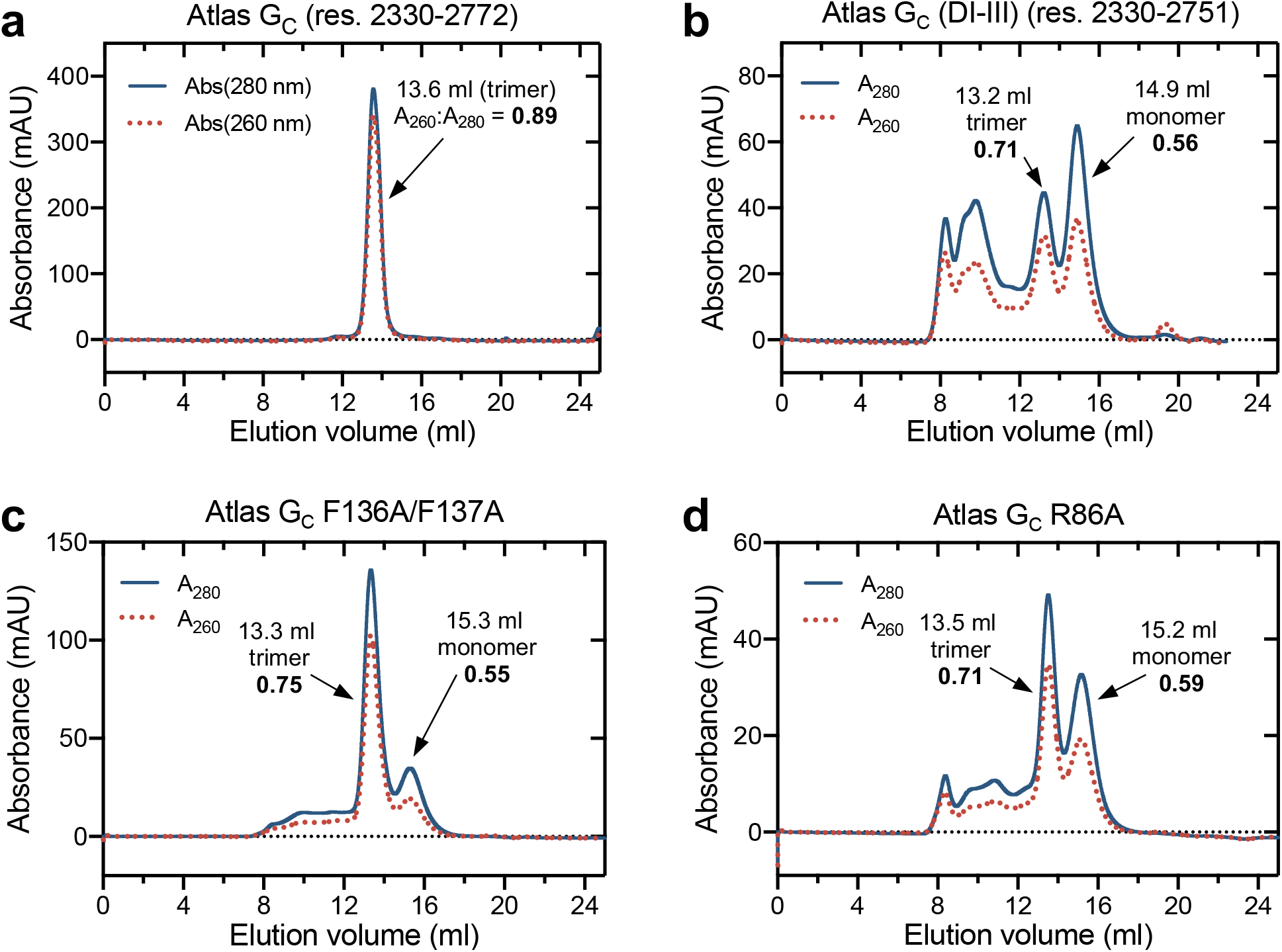
Size-exclusion chromatography of Atlas G_C_ following ion-exchange chromatography. Samples were analysed on a Superdex 200 Increase (10/300) column (Cytiva). The A_260_:A_280_ ratios for selected peaks are shown in bold. **(a)** The Atlas G_C_ fragment used for cryo-EM image reconstruction (residues 2330-2772). The protein had higher absorbance at 260 nm (A_260_) than expected for pure protein. The elution volume was consistent with a homotrimer. **(b)** Atlas G_C_(DI-III), maintained at pH > 7 throughout expression and purification. Multiple oligomeric states were present including peaks with elution volumes consistent with monomers, trimers and higher order oligomers. **(c)** Atlas G_C_ fusion loop mutant F136A/F137A. **(d)** Atlas G_C_ GPL binding pocket mutant R86A.

**Supplementary Fig. 3.**
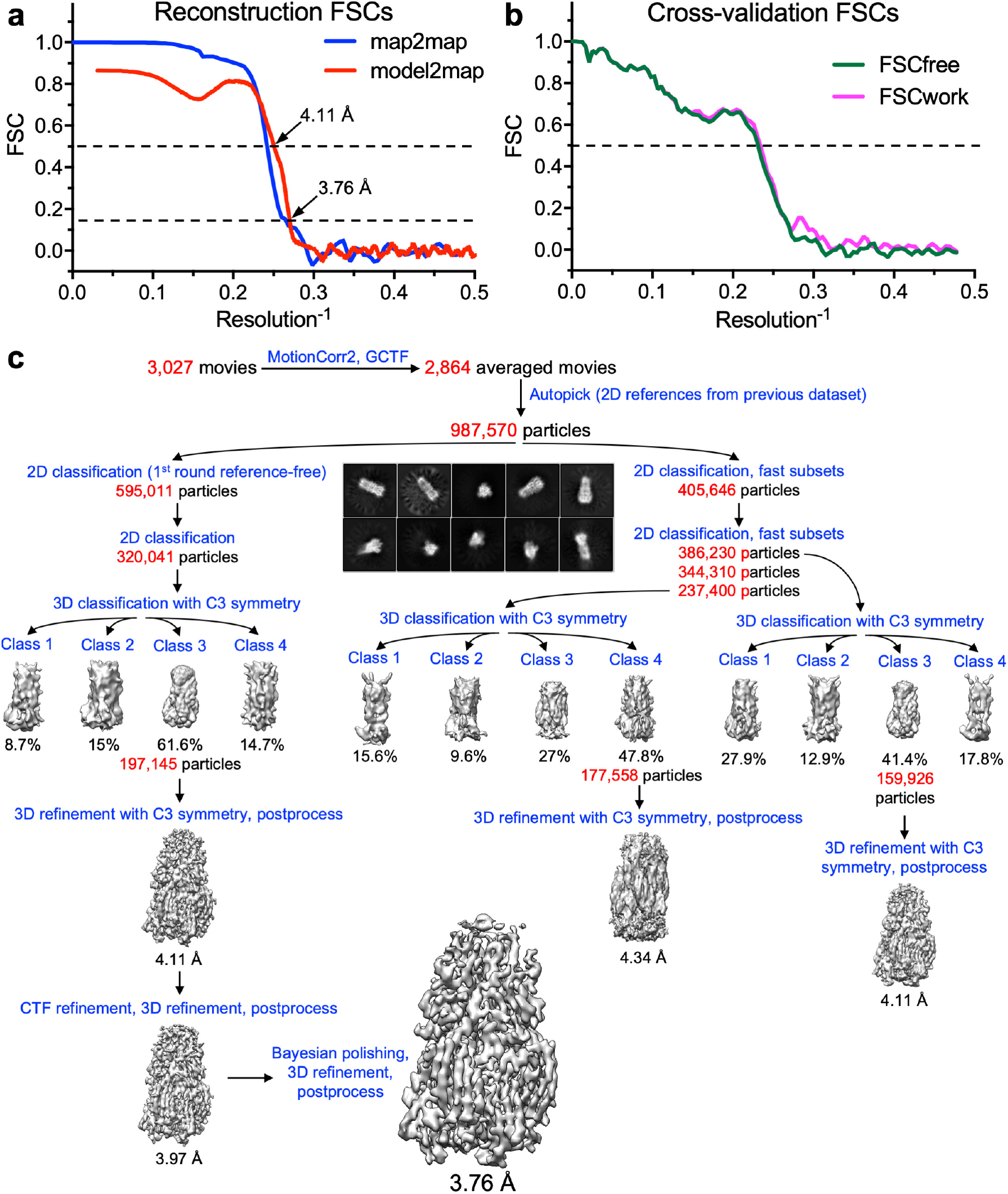
Image reconstruction quality assessment and workflow. (**a**) Fourier Shell Correlations (FSC) of the reconstruction of Atlas G_C_ from two independently refined half-maps (map2map, in blue); and of the reconstruction from the whole dataset versus a map calculated from the refined atomic model (model2map, in red). The gold-standard cutoff (FSC = 0.143) and the FSC = 0.5 level are marked with dashed lines. The resolution values of each curve at these levels are indicated. (**b**) FSC plots for cross-validation as described by Amunts *et al.* (*74*). FSCwork (magenta), FSC of refined test model versus work set (half-map used in test refinement). FSCfree (green), FSC of refined test model versus test set (half-map not used in test refinement). The FSC = 0.5 level is indicated by a dashed line. (**c**) Flow chart showing the workflow pipeline for cryo-EM image processing, classification and model refinement, as described in the Materials and Methods.

**Supplementary Fig. 4.**
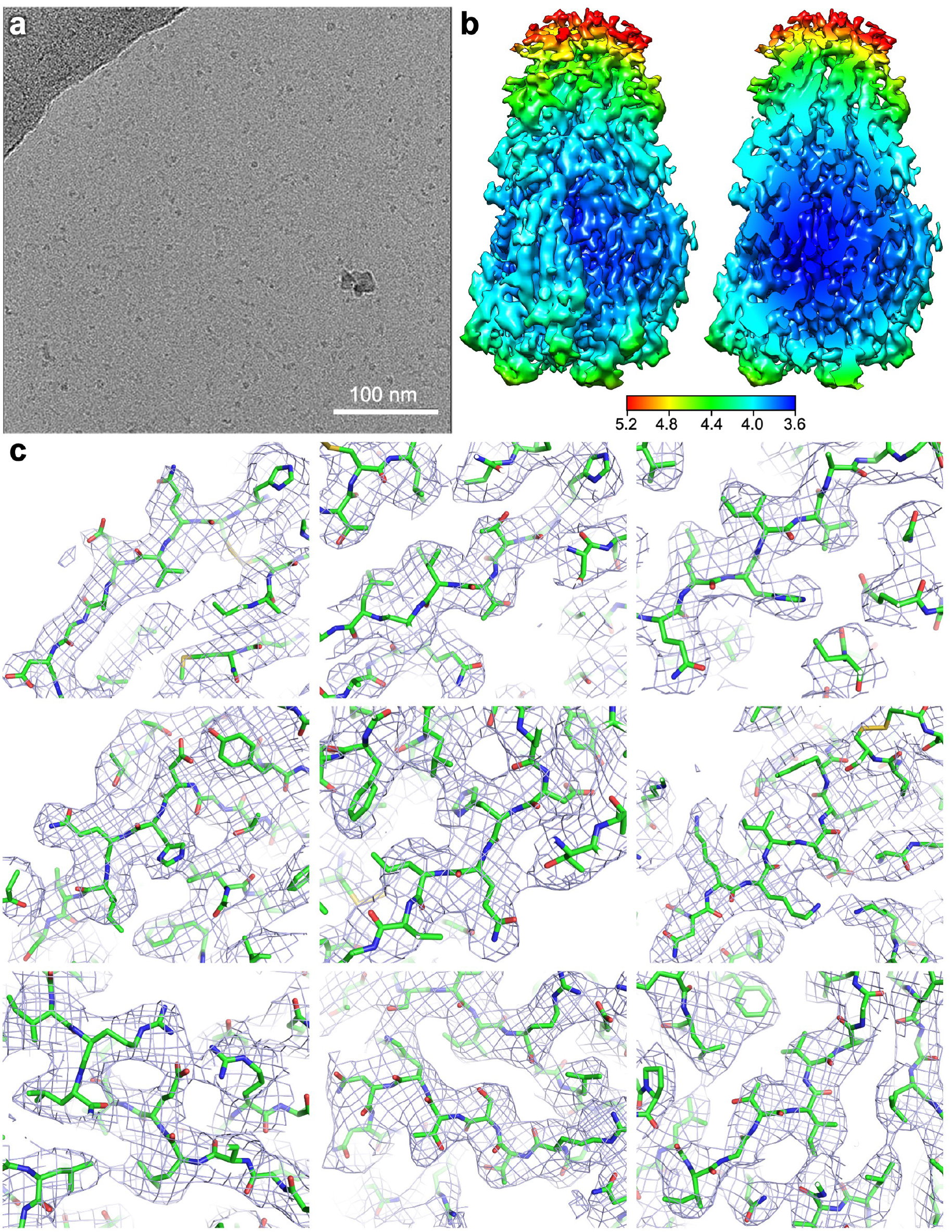
Representative Cryo-EM micrograph and density with local resolution estimation. (**a**) Representative cryo-EM micrograph of Atlas G_C_ trimers. (**b**) Local resolution estimation for the cryo-EM volume, calculated in RELION 3.0 (*69*). (**c**) Representative samples of local cryo-EM density from the structure with fitted and refined atomic models. The deposited density map was contoured at 3 *σ* in Pymol (Schrodinger, LLC).

**Supplementary Fig. 5.**
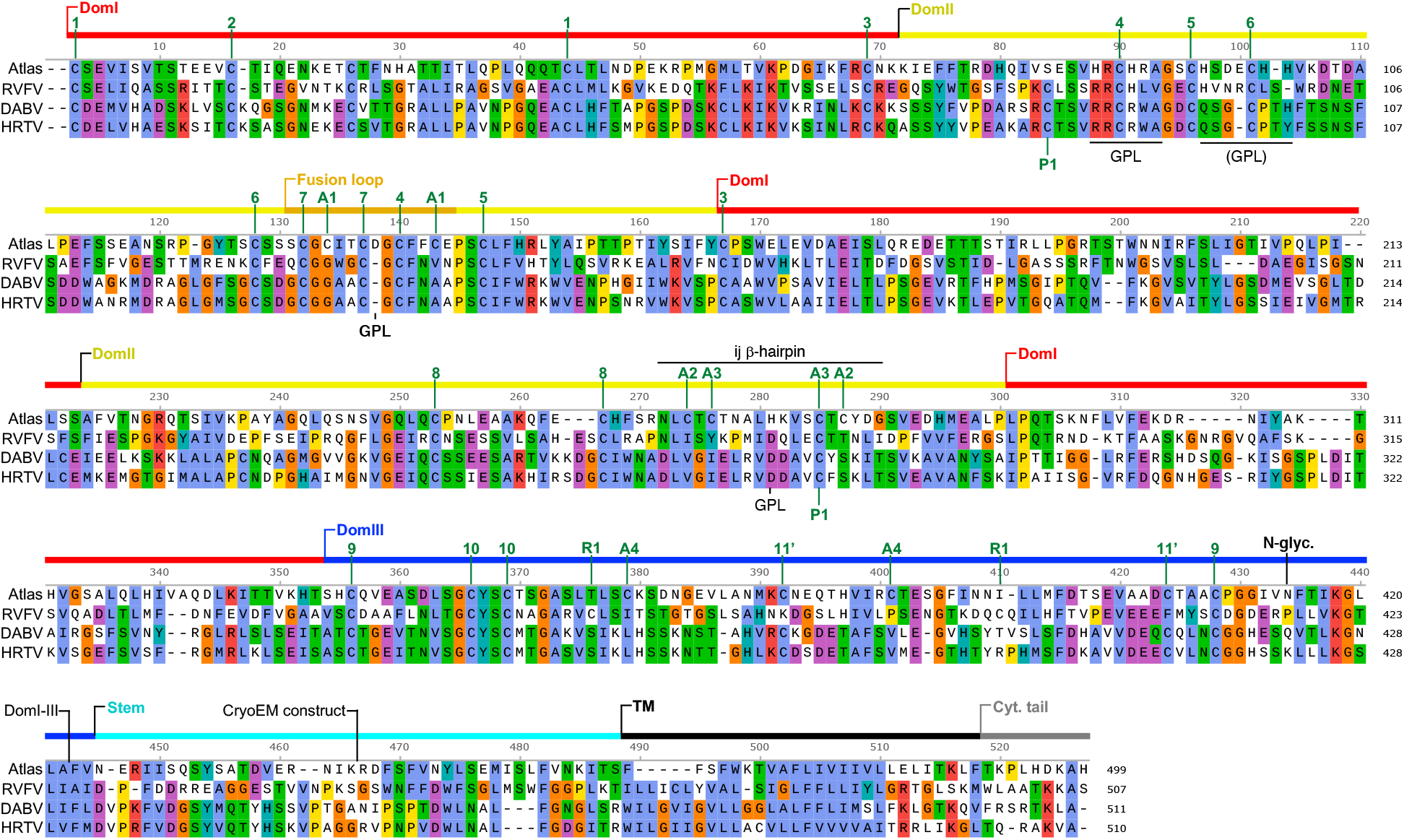
Protein sequence alignment of Atlas G_C_ and its closest orthologs. Amino acid sequence alignment of Atlas G_C_ and the most similar sequences from infectious viruses: Rift Valley Fever virus (RVFV), Dabie bandavirus (DABV, formerly SFTSV phlebovirus) and Heartland virus (HRTV). Domains and structural features are marked with lines above the alignment. GPL, residues in or near the glycerophospholipid headgroup binding pocket. N-glyc., N-glycosylation site. Cyt. tail, cytoplasmic tail. Conserved disulfide bonds are numbered in green. Disulfide bonds specific to Atlas virus are denoted with an “A”. An RVFV-specific disulfide is denoted with an “R”. A phlebovirus-specific disulfide is denoted with a “P”. The disulfide marked 11’ is conserved in Atlas, DABV and HRTV but not RVFV.

**Supplementary Fig. 6.**
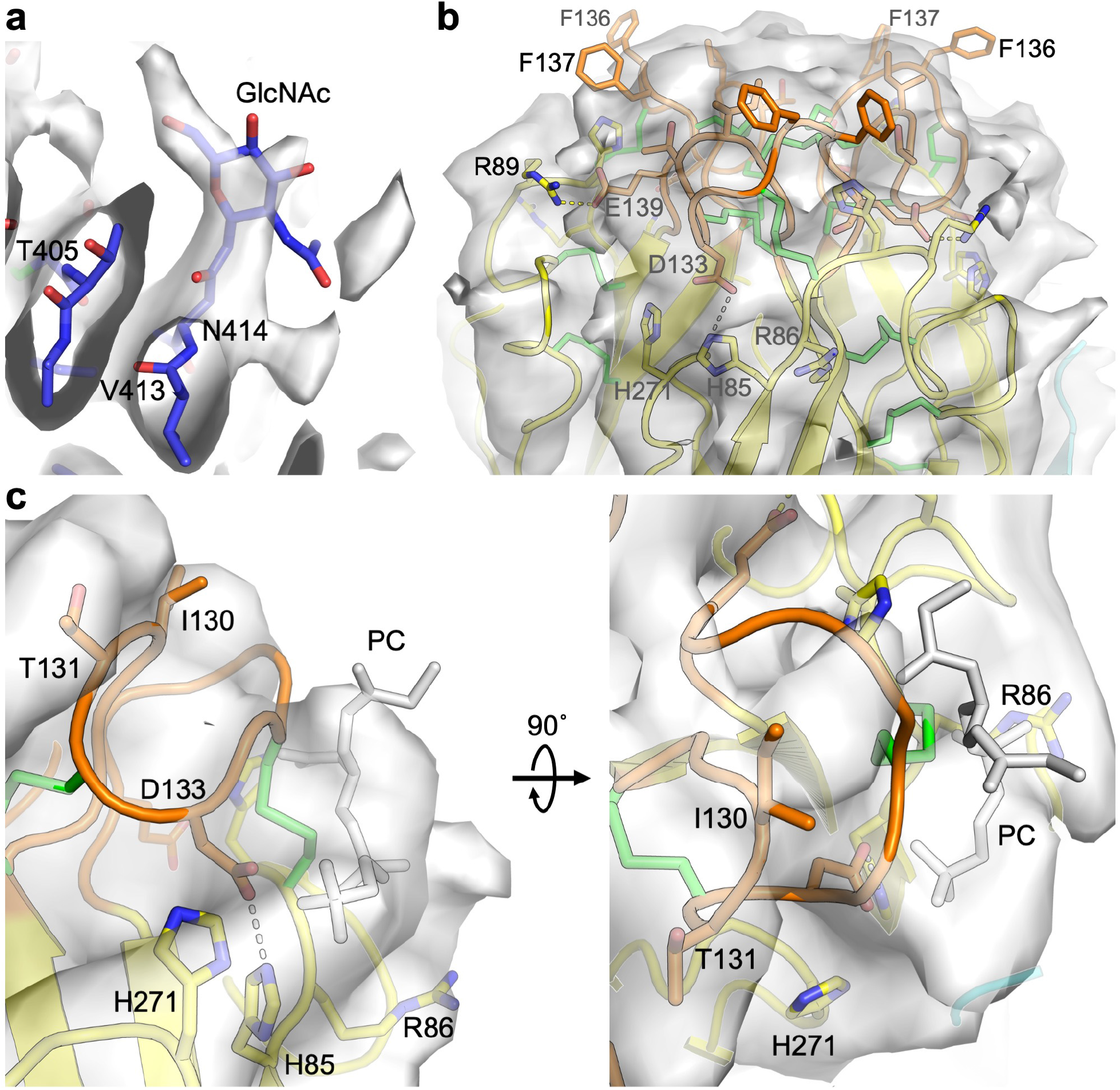
Cryo-EM density for N-linked glycan and GPL pocket of Atlas G_C_. **(a)** Cryo-EM density and atomic model of the N-linked glycan at Asn414. The deposited density map was contoured at 2.2 *σ* in Pymol (Schrodinger, LLC). **(b)** Cryo-EM density and atomic model for the fusion loop region of the Atlas G_C_ trimer. A B-factor blurring correction of +150 Å^2^ was applied to the deposited density map. The resulting map was contoured at 1 *σ* in Pymol. **(c)** Cryo-EM density in the glycerophospholipid (GPL) headgroup binding pocket unaccounted for by the atomic model. Shown in grey stick representation is the phosphatidylcholine (PC) headgroup bound to RVFV G_C_ structure (*27*) superimposed on the Atlas G_C_ structure. A B-factor blurring correction of +163 Å^2^ was applied to the deposited density map. The resulting map was contoured at 1.2 *σ* in Pymol.

**Supplementary Fig. 7.**
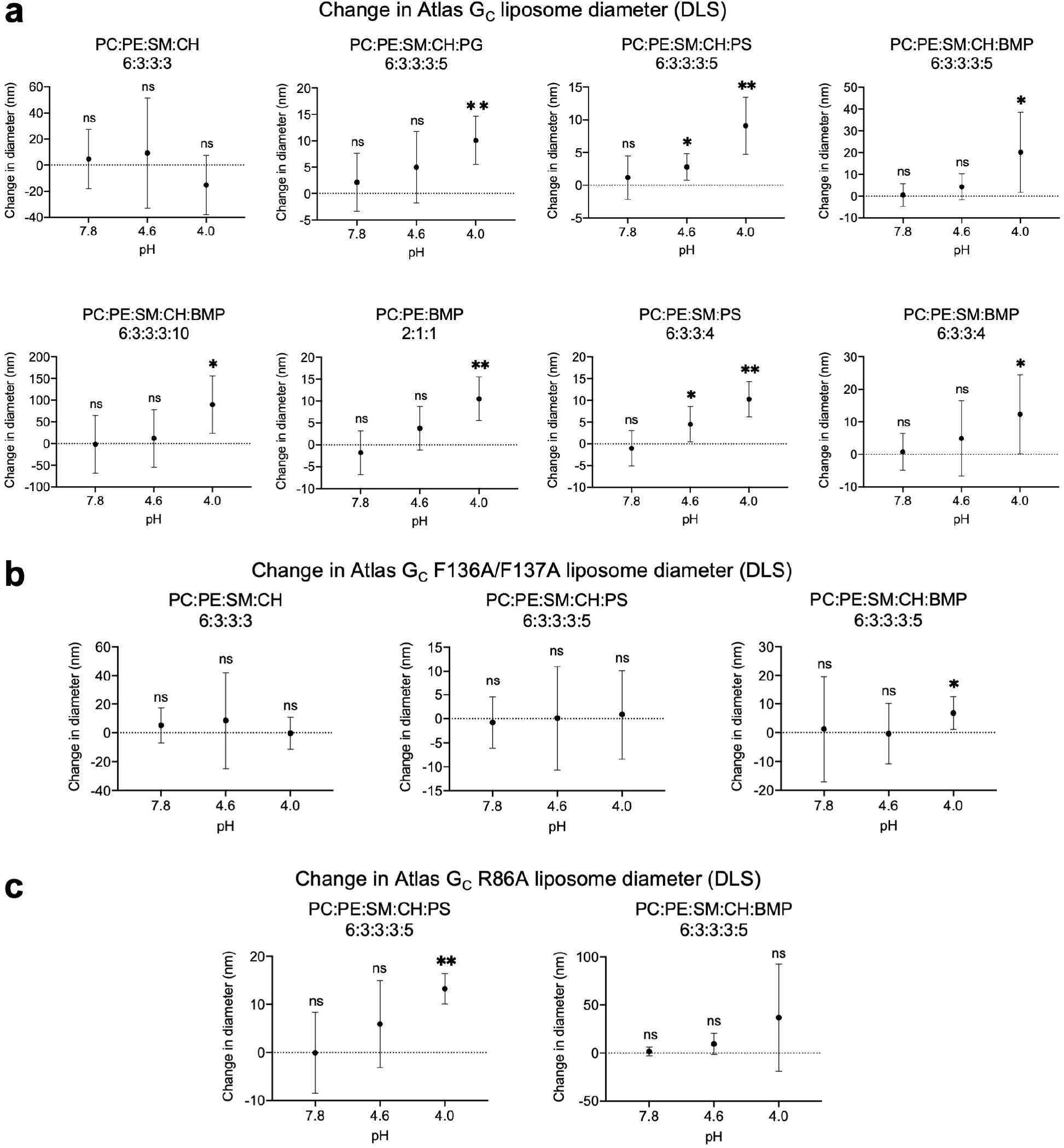
Dynamic light scattering of Atlas G_C_-liposome complexes. Liposome diameter dynamic light scattering (DLS) data from Fig. 4 are shown as the mean change in liposome diameter. **(a)** Change in liposome diameter upon binding Atlas G_C_ ectodomain. **(b)** Change in liposome diameter upon binding Atlas G_C_ F136A/F137A. **(c)** Change in liposome diameter upon binding Atlas G_C_ R86A. Error bars show the standard error of the mean (s.e.m.) of three to seven replicates (see **Supplementary Dataset 1** for source data). Significance was determined by 2-way ANOVA analysis using Sidak’s multiple comparisons test with a 95% confidence interval, in Prism 8 (GraphPad). **, 0.001 < *p* < 0.01; *, *p* < 0.05; ns, not significant.

**Supplementary Fig. 8.**
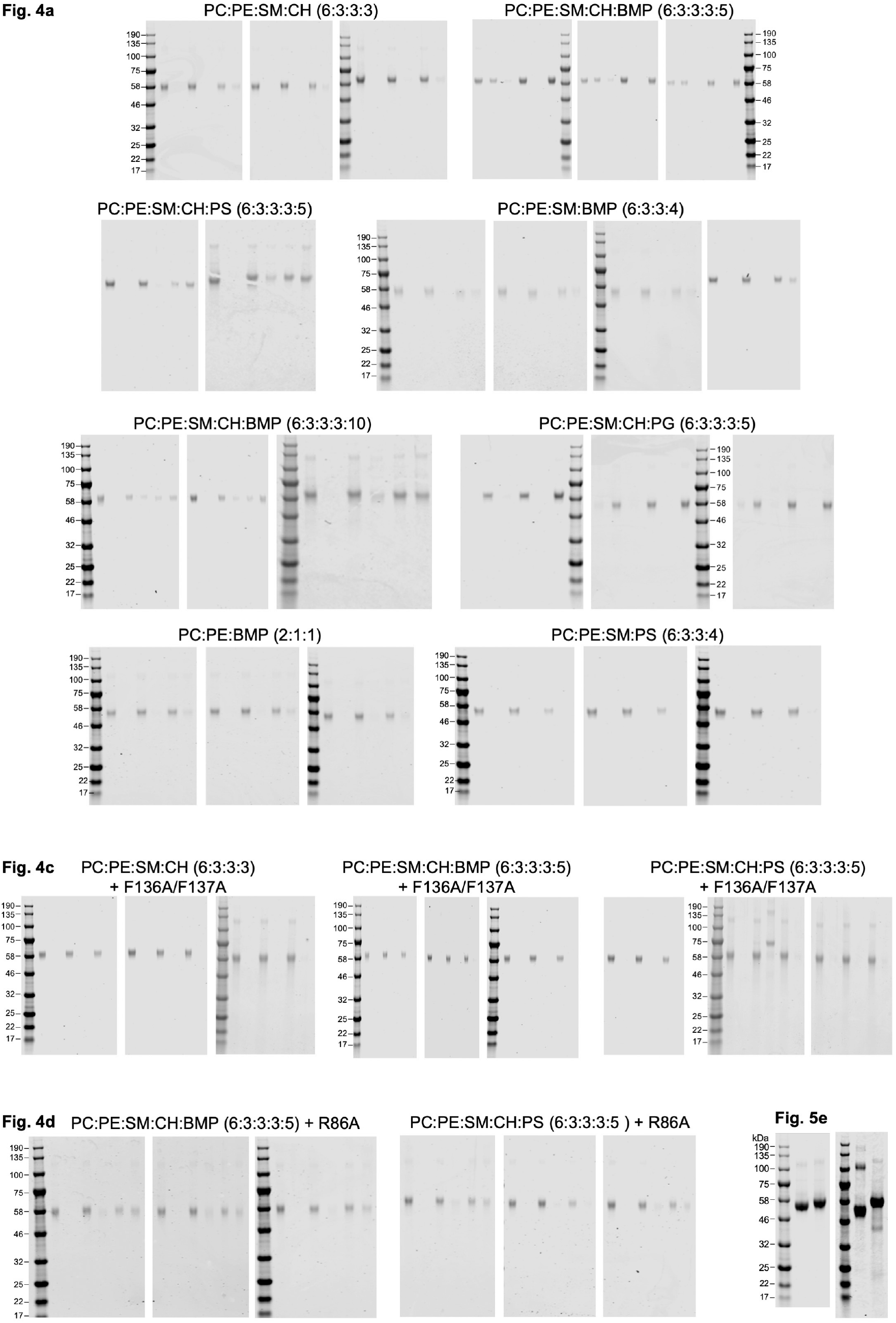
Uncropped gels from Figures 4 and 5. Gels are shown for all liposome flotation assay replicates (Fig. 4) and non-reducing gels (Fig. 5).

**Supplementary Fig. 9.**
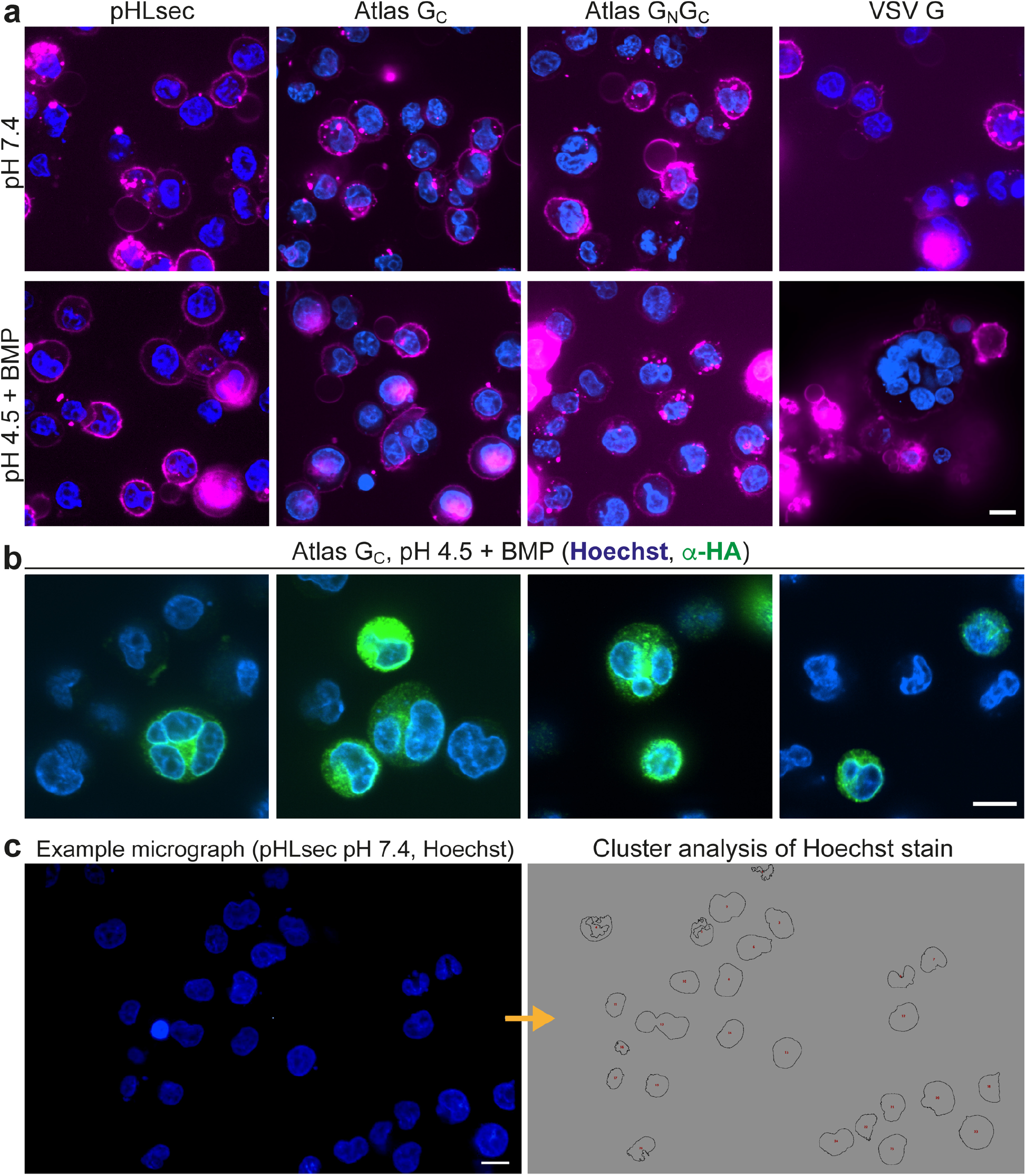
Cell-cell fusion assay with CHO cells. **(a)** Representative confocal micrographs of CHO cells transfected with plasmid encoding Atlas G_C_, Atlas G_N_-G_C_, VSV G, or no protein (pHLsec empty vector). Cells were treated with pH 4.5 buffer containing BMP, or pH 7.4 buffer without exogenous lipid. Blue, Hoechst 33342 nuclear stain; Magenta, CellBrite Red plasma membrane dye. **(b)** Permeabilized and fixed cells transfected with plasmid encoding G_C_ stained with anti-HA antibody, to detect the C-terminal HA tag on G_C_. Blue, Hoechst 33342; green, anti-HA antibody. **(c)** Left, confocal micrograph of cells transfected with pHLsec plasmid stained with Hoechst 33342. Right, masks from cluster analysis of the Hoechst channel with Fiji (*76*). All scale bars in this figure are 10 µm.

**Supplementary Fig. 10.**
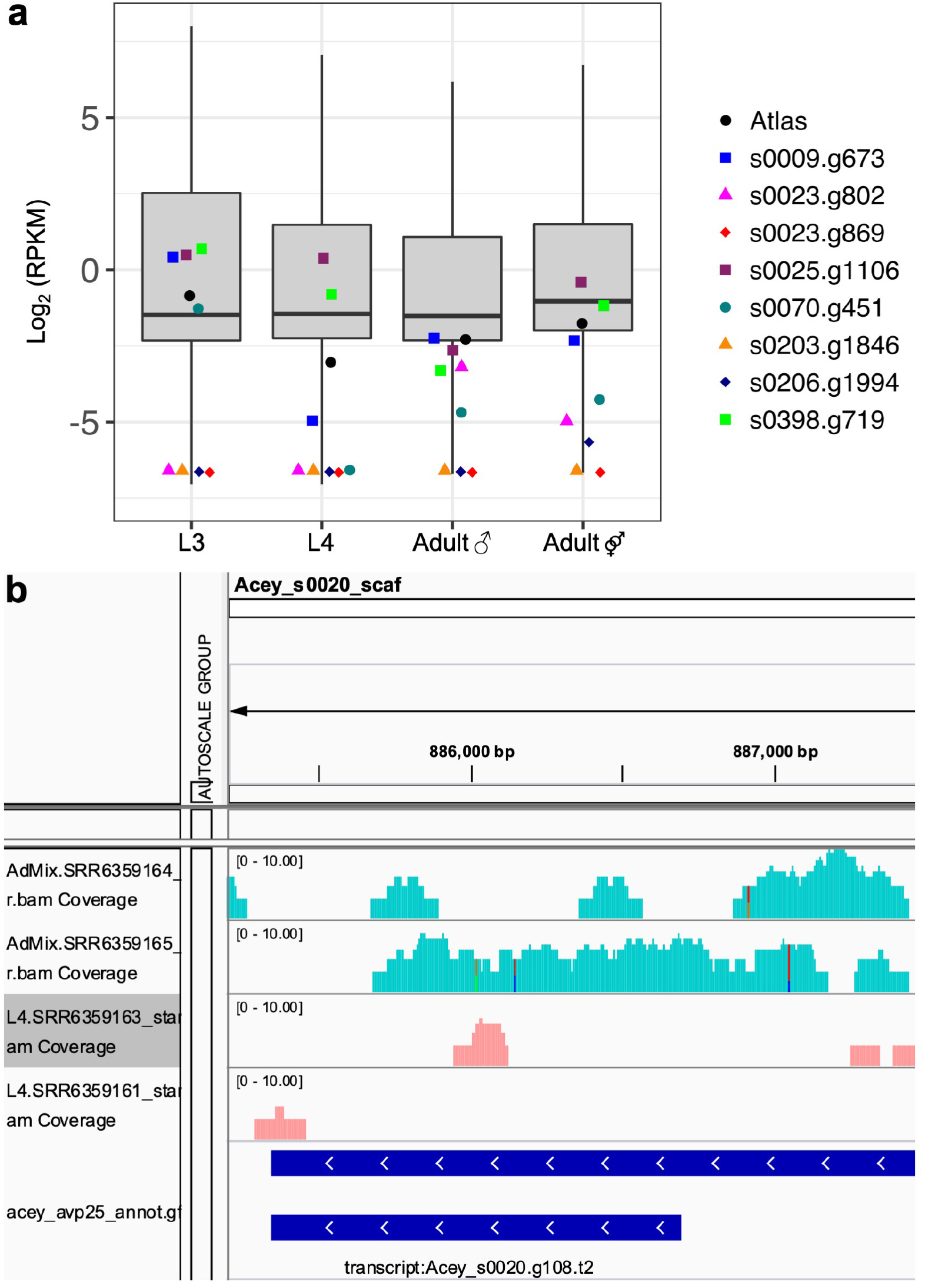
RNA-Seq analysis of intact *A. ceylanicum* belpaovirus ERVs. **(a)** Boxplot representation of RNA-Seq expression in Reads Per Kilobase of transcript per Million mapped reads (RPKM) for all annotated genes, for Atlas virus, and for the eight other intact *A. ceylanicum* belpaovirus ERVs at the L3, L4 and adult (male and mixed sex) developmental stages. RNA-Seq data are from Bernot *et al.* (*65*). **(b)** Integrative Genomics Viewer (IGV) snapshot showing the distribution of raw RNA-Seq reads over the Atlas virus (*Acey_s0020.g108*) coding sequence in mixed sex adult and L4 worms with individual unmerged BAM files.

**Supplementary Table 1.**
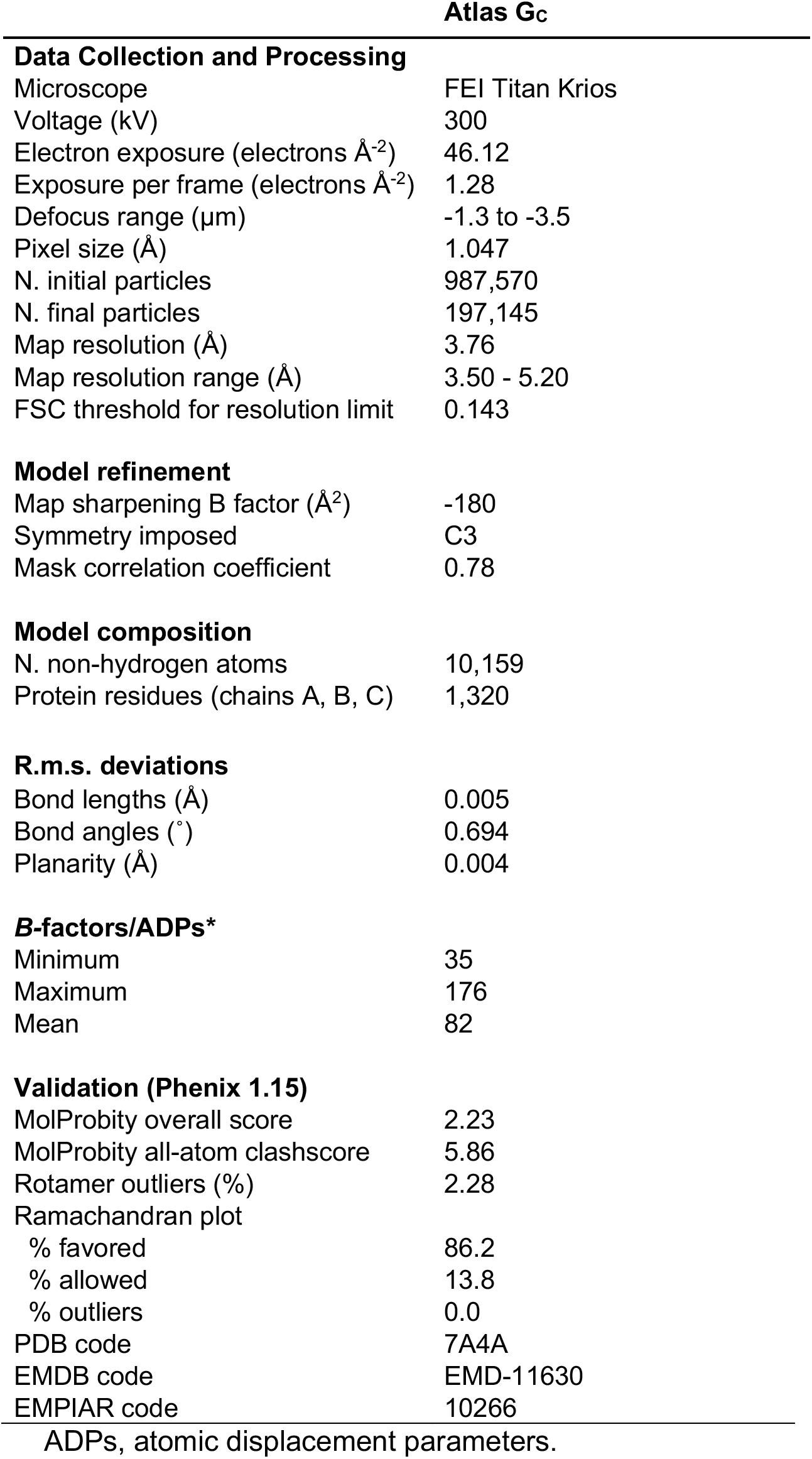
Cryo-EM data collection, structure determination, model building and refinement parameters and statistics.

## References

1. M. Friedli, D. Trono, The developmental control of transposable elements and the evolution of higher species. Annu Rev Cell Dev Biol 31, 429–451 (2015).

2. E. B. Chuong, N. C. Elde, C. Feschotte, Regulatory evolution of innate immunity through co-option of endogenous retroviruses. Science 351, 1083–1087 (2016).

3. J. A. Frank, C. Feschotte, Co-option of endogenous viral sequences for host cell function. Curr Opin Virol 25, 81–89 (2017).

4. A. Dupressoir, C. Lavialle, T. Heidmann, From ancestral infectious retroviruses to bona fide cellular genes: role of the captured syncytins in placentation. Placenta 33, 663–671 (2012).

5. R. S. Moore et al., Horizontal and vertical transmission of transgenerational memories via the *Cer1* transposon. bioRxiv, 2020.2012.2028.424563 (2020).

6. T. Gojobori, S. Yokoyama, Rates of evolution of the retroviral oncogene of Moloney murine sarcoma virus and of its cellular homologues. Proc Natl Acad Sci U S A 82, 4198–4201 (1985).

7. K. R. McCarthy et al., The ancient fusogen EnvP(b)1 is expressed in human tissues and its structure informs the evolution of gammaretrovirus envelope proteins. bioRxiv, 2020.2004.2022.056234 (2020).

8. D. Kremer et al., Human endogenous retrovirus type W envelope protein inhibits oligodendroglial precursor cell differentiation. Ann Neurol 74, 721–732 (2013).

9. W. Li et al., Human endogenous retrovirus-K contributes to motor neuron disease. Sci Transl Med 7, 307ra153 (2015).

10. M. Krupovic et al., Ortervirales: New Virus Order Unifying Five Families of Reverse-Transcribing Viruses. J Virol 92, (2018).

11. I. G. Frame, J. F. Cutfield, R. T. Poulter, New BEL-like LTR-retrotransposons in Fugu rubripes, Caenorhabditis elegans, and Drosophila melanogaster. Gene 263, 219–230 (2001).

12. N. J. Bowen, J. F. McDonald, Genomic analysis of Caenorhabditis elegans reveals ancient families of retroviral-like elements. Genome Res 9, 924–935 (1999).

13. H. S. Malik, S. Henikoff, T. H. Eickbush, Poised for contagion: evolutionary origins of the infectious abilities of invertebrate retroviruses. Genome Res 10, 1307–1318 (2000).

14. D. Fass, S. C. Harrison, P. S. Kim, Retrovirus envelope domain at 1.7 angstrom resolution. Nat Struct Biol 3, 465–469 (1996).

15. D. C. Chan, D. Fass, J. M. Berger, P. S. Kim, Core structure of gp41 from the HIV envelope glycoprotein. Cell 89, 263–273 (1997).

16. W. Weissenhorn, A. Dessen, S. C. Harrison, J. J. Skehel, D. C. Wiley, Atomic structure of the ectodomain from HIV-1 gp41. Nature 387, 426–430 (1997).

17. B. Kobe, R. J. Center, B. E. Kemp, P. Poumbourios, Crystal structure of human T cell leukemia virus type 1 gp21 ectodomain crystallized as a maltose-binding protein chimera reveals structural evolution of retroviral transmembrane proteins. Proc Natl Acad Sci U S A 96, 4319–4324 (1999).

18. S. C. Harrison, Viral membrane fusion. Virology479-480, 498–507 (2015).

19. M. Dessau, Y. Modis, Crystal structure of glycoprotein C from Rift Valley fever virus. Proc. Natl. Acad. Sci. U. S. A. 110, 1696–1701 (2013).

20. Y. Modis, Relating Structure to Evolution in Class II Viral Membrane Fusion Proteins. Curr Opin Virol, DOI:10.1016/j.coviro.2014.1001.1009 (2014).

21. J. Lescar et al., The Fusion glycoprotein shell of Semliki Forest virus: an icosahedral assembly primed for fusogenic activation at endosomal pH. Cell 105, 137–148 (2001).

22. F. A. Rey, F. X. Heinz, C. Mandl, C. Kunz, S. C. Harrison, The envelope glycoprotein from tick-borne encephalitis virus at 2 A resolution. Nature 375, 291–298 (1995).

23. Y. Modis, S. Ogata, D. Clements, S. C. Harrison, A ligand-binding pocket in the dengue virus envelope glycoprotein. Proc Natl Acad Sci U S A 100, 6986–6991 (2003).

24. R. M. DuBois et al., Functional and evolutionary insight from the crystal structure of rubella virus protein E1. Nature 493, 552–556 (2013).

25. Y. Modis, S. Ogata, D. Clements, S. C. Harrison, Structure of the dengue virus envelope protein after membrane fusion. Nature 427, 313–319 (2004).

26. D. L. Gibbons et al., Conformational change and protein-protein interactions of the fusion protein of Semliki Forest virus. Nature 427, 320–325 (2004).

27. P. Guardado-Calvo et al., A glycerophospholipid-specific pocket in the RVFV class II fusion protein drives target membrane insertion. Science 358, 663–667 (2017).

28. J. Perez-Vargas et al., Structural basis of eukaryotic cell-cell fusion. Cell 157, 407–419 (2014).

29. J. Fedry et al., The Ancient Gamete Fusogen HAP2 Is a Eukaryotic Class II Fusion Protein. Cell 168, 904–915 e910 (2017).

30. J. Fedry et al., Evolutionary diversification of the HAP2 membrane insertion motifs to drive gamete fusion across eukaryotes. PLoS Biol 16, e2006357 (2018).

31. J. Feng et al., Fusion surface structure, function, and dynamics of gamete fusogen HAP2. eLife 7, (2018).

32. T. Clark, HAP2/GCS1: Mounting evidence of our true biological EVE? PLoS Biol 16, e3000007 (2018).

33. S. F. Altschul et al., Gapped BLAST and PSI-BLAST: a new generation of protein database search programs. Nucleic Acids Res 25, 3389–3402 (1997).

34. O. Poch, I. Sauvaget, M. Delarue, N. Tordo, Identification of four conserved motifs among the RNA-dependent polymerase encoding elements. EMBO J 8, 3867–3874 (1989).

35. H. Felder et al., Tas, a retrotransposon from the parasitic nematode Ascaris lumbricoides. Gene 149, 219–225 (1994).

36. I. Letunic, P. Bork, Interactive Tree Of Life (iTOL) v4: recent updates and new developments. Nucleic Acids Res 47, W256–W259 (2019).

37. S. Halldorsson et al., Structure of a phleboviral envelope glycoprotein reveals a consolidated model of membrane fusion. Proc Natl Acad Sci U S A 113, 7154–7159 (2016).

38. Y. Zhu et al., The Postfusion Structure of the Heartland Virus Gc Glycoprotein Supports Taxonomic Separation of the Bunyaviral Families Phenuiviridae and Hantaviridae. J Virol 92, (2018).

39. D. J. Leahy, W. A. Hendrickson, I. Aukhil, H. P. Erickson, Structure of a fibronectin type III domain from tenascin phased by MAD analysis of the selenomethionyl protein. Science 258, 987–991 (1992).

40. J. F. Pinello et al., Structure-Function Studies Link Class II Viral Fusogens with the Ancestral Gamete Fusion Protein HAP2. Curr Biol 27, 651–660 (2017).

41. M. Kielian, F. A. Rey, Virus membrane-fusion proteins: more than one way to make a hairpin. Nat. Rev. Microbiol. 4, 67–76 (2006).

42. L. Holm, P. Rosenstrom, Dali server: conservation mapping in 3D. Nucleic Acids Res 38, W545–549 (2010).

43. D. E. Klein, J. L. Choi, S. C. Harrison, Structure of a dengue virus envelope protein late-stage fusion intermediate. J Virol 87, 2287–2293 (2013).

44. S. Murakami, K. Terasaki, S. I. Ramirez, J. C. Morrill, S. Makino, Development of a novel, single-cycle replicable rift valley Fever vaccine. PLoS neglected tropical diseases 8, e2746 (2014).

45. D. Bitto, S. Halldorsson, A. Caputo, J. T. Huiskonen, Low pH and Anionic Lipid-dependent Fusion of Uukuniemi Phlebovirus to Liposomes. J Biol Chem 291, 6412–6422 (2016).

46. E. Zaitseva, S. T. Yang, K. Melikov, S. Pourmal, L. V. Chernomordik, Dengue virus ensures its fusion in late endosomes using compartment-specific lipids. PLOS Pathog. 6, e1001131 (2010).

47. D. L. Esposito, J. B. Nguyen, D. C. DeWitt, E. Rhoades, Y. Modis, Physico-chemical requirements and kinetics of membrane fusion of flavivirus-like particles. J Gen Virol 96, 1702–1711 (2015).

48. A. M. Nour, Y. Li, J. Wolenski, Y. Modis, Viral membrane fusion and nucleocapsid delivery into the cytoplasm are distinct events in some flaviviruses. PLoS Pathog 9, e1003585 (2013).

49. P. Y. Lozach et al., Entry of bunyaviruses into mammalian cells. Cell Host Microbe 7, 488–499 (2010).

50. M. Umashankar et al., Differential cholesterol binding by class II fusion proteins determines membrane fusion properties. J Virol 82, 9245–9253 (2008).

51. P. K. Chatterjee, C. H. Eng, M. Kielian, Novel mutations that control the sphingolipid and cholesterol dependence of the Semliki Forest virus fusion protein. J Virol 76, 12712–12722 (2002).

52. J. M. Smit, R. Bittman, J. Wilschut, Low-pH-dependent fusion of Sindbis virus with receptor-free cholesterol- and sphingolipid-containing liposomes. J Virol 73, 8476–8484 (1999).

53. T. V. Kurzchalia, S. Ward, Why do worms need cholesterol? Nat Cell Biol 5, 684–688 (2003).

54. M. Merris, J. Kraeft, G. S. Tint, J. Lenard, Long-term effects of sterol depletion in C. elegans: sterol content of synchronized wild-type and mutant populations. J Lipid Res 45, 2044–2051 (2004).

55. M. Ruiz et al., Membrane fluidity is regulated by the C. elegans transmembrane protein FLD-1 and its human homologs TLCD1/2. eLife 7, (2018).

56. K. Stiasny, S. L. Allison, J. Schalich, F. X. Heinz, Membrane interactions of the tick-borne encephalitis virus fusion protein E at low pH. J. Virol. 76, 3784–3790 (2002).

57. D. L. Gibbons, A. Ahn, P. K. Chatterjee, M. Kielian, Formation and characterization of the trimeric form of the fusion protein of Semliki Forest Virus. J Virol 74, 7772–7780 (2000).

58. R. Fritz, K. Stiasny, F. X. Heinz, Identification of specific histidines as pH sensors in flavivirus membrane fusion. J Cell Biol 183, 353–361 (2008).

59. V. Nayak et al., Crystal structure of dengue virus type 1 envelope protein in the postfusion conformation and its implications for membrane fusion. J. Virol. 83, 4338–4344 (2009).

60. Z. L. Qin, Y. Zheng, M. Kielian, Role of conserved histidine residues in the low-pH dependence of the Semliki Forest virus fusion protein. J Virol 83, 4670–4677 (2009).

61. Y. Zheng, C. Sanchez-San Martin, Z. L. Qin, M. Kielian, The domain I-domain III linker plays an important role in the fusogenic conformational change of the alphavirus membrane fusion protein. J Virol 85, 6334–6342 (2011).

62. S. M. de Boer et al., Acid-activated structural reorganization of the Rift Valley fever virus Gc fusion protein. J Virol 86, 13642–13652 (2012).

63. M. A. Vega, J. L. Strominger, Constitutive endocytosis of HLA class I antigens requires a specific portion of the intracytoplasmic tail that shares structural features with other endocytosed molecules. Proc Natl Acad Sci U S A 86, 2688–2692 (1989).

64. R. DeMarco et al., Saci-1, -2, and -3 and Perere, four novel retrotransposons with high transcriptional activities from the human parasite Schistosoma mansoni. J Virol 78, 2967–2978 (2004).

65. J. P. Bernot et al., Transcriptomic analysis of hookworm Ancylostoma ceylanicum life cycle stages reveals changes in G-protein coupled receptor diversity associated with the onset of parasitism. Int J Parasitol 50, 603–610 (2020).

66. S. Q. Zheng et al., MotionCor2: anisotropic correction of beam-induced motion for improved cryo-electron microscopy. Nat Methods 14, 331–332 (2017).

67. K. Zhang, Gctf: Real-time CTF determination and correction. J Struct Biol 193, 1–12 (2016).

68. S. He, S. H. W. Scheres, Helical reconstruction in RELION. J Struct Biol 198, 163–176 (2017).

69. S. H. Scheres, RELION: implementation of a Bayesian approach to cryo-EM structure determination. J Struct Biol 180, 519–530 (2012).

70. J. Zivanov, T. Nakane, S. H. W. Scheres, A Bayesian approach to beam-induced motion correction in cryo-EM single-particle analysis. IUCrJ 6, 5–17 (2019).

71. E. F. Pettersen et al., UCSF Chimera--a visualization system for exploratory research and analysis. J Comput Chem 25, 1605–1612 (2004).

72. P. D. Adams et al., PHENIX: a comprehensive Python-based system for macromolecular structure solution. Acta Crystallogr D Biol Crystallogr 66, 213–221 (2010).

73. P. Emsley, K. Cowtan, Coot: model-building tools for molecular graphics. Acta Crystallogr D Biol Crystallogr 60, 2126–2132 (2004).

74. A. Amunts et al., Structure of the yeast mitochondrial large ribosomal subunit. Science 343, 1485–1489 (2014).

75. V. B. Chen et al., MolProbity: all-atom structure validation for macromolecular crystallography. Acta Crystallogr D Biol Crystallogr 66, 12–21 (2010).

76. J. Schindelin et al., Fiji: an open-source platform for biological-image analysis. Nat Methods 9, 676–682 (2012).

77. A. Teissandier, N. Servant, E. Barillot, D. Bourc’his, Tools and best practices for retrotransposon analysis using high-throughput sequencing data. Mob DNA 10, 52 (2019).

78. E. M. Schwarz et al., The genome and transcriptome of the zoonotic hookworm Ancylostoma ceylanicum identify infection-specific gene families. Nat Genet 47, 416–422 (2015).

79. A. Dobin, T. R. Gingeras, Mapping RNA-seq Reads with STAR. Curr Protoc Bioinformatics 51, 11 14 11–11 14 19 (2015).

80. Y. Liao, G. K. Smyth, W. Shi, featureCounts: an efficient general purpose program for assigning sequence reads to genomic features. Bioinformatics 30, 923–930 (2014).

81. K. L. Howe, B. J. Bolt, M. Shafie, P. Kersey, M. Berriman, WormBase ParaSite - a comprehensive resource for helminth genomics. Molecular and biochemical parasitology 215, 2–10 (2017).

82. R. C. Team. (R Foundation for Statistical Computing, Vienna, Austria, 2019).

83. M. I. Love, W. Huber, S. Anders, Moderated estimation of fold change and dispersion for RNA-seq data with DESeq2. Genome Biol 15, 550 (2014).

84. H. Wickham, ggplot2: Elegant Graphics for Data Analysis. (Springer-Verlag, New York, 2016).

85. B. Gel, E. Serra, karyoploteR: an R/Bioconductor package to plot customizable genomes displaying arbitrary data. Bioinformatics 33, 3088–3090 (2017).

86. S. Kurtz et al., REPuter: the manifold applications of repeat analysis on a genomic scale. Nucleic Acids Res 29, 4633–4642 (2001).

87. M. Krupovic, E. V. Koonin, Homologous Capsid Proteins Testify to the Common Ancestry of Retroviruses, Caulimoviruses, Pseudoviruses, and Metaviruses. J Virol 91, (2017).

88. B. M. Muhire, A. Varsani, D. P. Martin, SDT: a virus classification tool based on pairwise sequence alignment and identity calculation. PLoS One 9, e108277 (2014).

